# qgg: an R package for large-scale quantitative genetic analyses

**DOI:** 10.1101/503631

**Authors:** Palle Duun Rohde, Izel Fourie Sørensen, Peter Sørensen

**Affiliations:** Centre for Quantitative Genetics and Genomics, Department of Molecular Biology and Genetics, Aarhus University, 8830 Tjele, Denmark

## Abstract

**Summary:** Here, we present the R package **qgg**, which provides an environment for large-scale genetic analyses of quantitative traits and diseases. The qgg package provides an infrastructure for efficient processing of large-scale genetic data and functions for estimating genetic parameters, performing single and multiple marker association analyses, and genomic-based predictions of phenotypes.

**Availability and implementation:** The R package qgg is freely available. For latest updates, user guides and example scripts, consult the main page http://psoerensen.github.io/qgg.

**Supplementary information:** Supplementary material is available.

## Introduction

Understanding the genetic basis of complex traits and diseases, which is an important topic across scientific disciplines, requires large-scale genotype-phenotype data. Functional genomics is accumulating data about DNA function at gene, RNA transcript and protein levels. By using knowledge from different biological layers we can improve the accuracy of genomic prediction, and provide biological insight into complex phenotypes.

Here, we present the R package **qgg**; a collection of statistical genetic methods for large-scale genetic analyses allowing leveraging of prior biological knowledge into the genetic analysis. Regions of the genome that links to different types of functional genomic information, such as genes, biological pathways, genetic interactions, gene expression patterns etc., are termed genomic features. Our hypothesis is that for any complex trait or disease, there exist genomic features that are enriched for causal genetic variants. Identifying the set of genomic features enriched for causal genetic variants provide insight into the genetic basis of the trait in question, and can therefore have important implications; for example, in understanding, treating, and preventing human diseases. Previously, we have demonstrated that integrating genomic features into the genetic analysis provides new biological knowledge of the genetic basis of complex traits and diseases across different species; fruit flies (Edwards *et al*., 2016; Rohde *et al*., 2018), dairy cattle (Edwards *et al*., 2015; Fang *et al*., 2018), pigs (Sarup *et al*., 2016) and humans (Rohde *et al*., 2016).

## Implementation and main functions

The qgg package is implemented in R (R Core Team, 2018) allowing users to utilise existing statistical and graphical facilities, and to develop efficient workflows utilising available genomic annotation resources, such as Bioconductor (Huber *et al*., 2015).

The qgg package uses a simple infrastructure with two main steps: 1) Prepare data for downstream analysis, which include preparing a genotype file, constructing genomic relationship matrices, and if necessary, prune genetic variants for linkage disequilibrium. 2) Perform the genetic analyses, which can include estimation of genetic parameters, single marker association analysis, gene set enrichment analysis, and prediction of phenotypes from genetic variants. Multi-core processing with openMP, multithreaded matrix operations implemented in BLAS libraries (OpenBLAS, ATLAS or MKL) and fast memory-efficient batch processing of genotype data stored in binary files (e.g. PLINK bedfiles, (Purcell *et al*., 2007)) allow the data processing and data analyses to be computationally efficient (see Supplementary Information, section 10).

The Supplementary Information contains details on the statistical models implemented in qgg, and example scripts using phenotypic and genetic data from the UK Biobank Resource (Bycroft *et al*., 2018). In addition, several tutorials using publicly available genotype and phenotype data (http://dgrp2.gnets.ncsu.edu/, Mackay *et al*. (2012)) are available at the qgg website http://psoerensen.github.io/qgg.

## Analysis of human height

To illustrate some of the core facilities of the qgg package, we performed a genetic analysis of human height using the White British cohort in the UK Biobank Resource (Bycroft *et al*., 2018). We excluded individuals that were related, had more than 5,000 missing genotypes or had chromosomal aneuploidy (*n* = 335, 744 observations). Single nucleotide polymorphisms (SNPs) with minor allele frequencies < 0.01 and SNPs located within the major histocompatibility complex were excluded from the analyses; yielding a total of 599,297 SNPs.

Using five training sets each of 50,000 individuals, we estimated the proportion of phenotypic variance explained by the SNPs to *ĥ*^2^= 0.69±0.004. We then partitioned the total genetic variance to variance captured by each autosomal chromosome (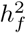, an example of a simple genomic feature, Fig. 1A). We computed polygenic risk scores for human height using SNP effects (pruned for linkage disequilibrium, *r*^2^< 0.7) from linear model association on different training-population sizes (100*K*, 200*K* and 300*K*). The maximum variance explained in the validation set was *R*^*2*^ = 0.22 at *p* < 0.05 for *n* = 300*K* (Fig. 1B).

**Figure 1.**
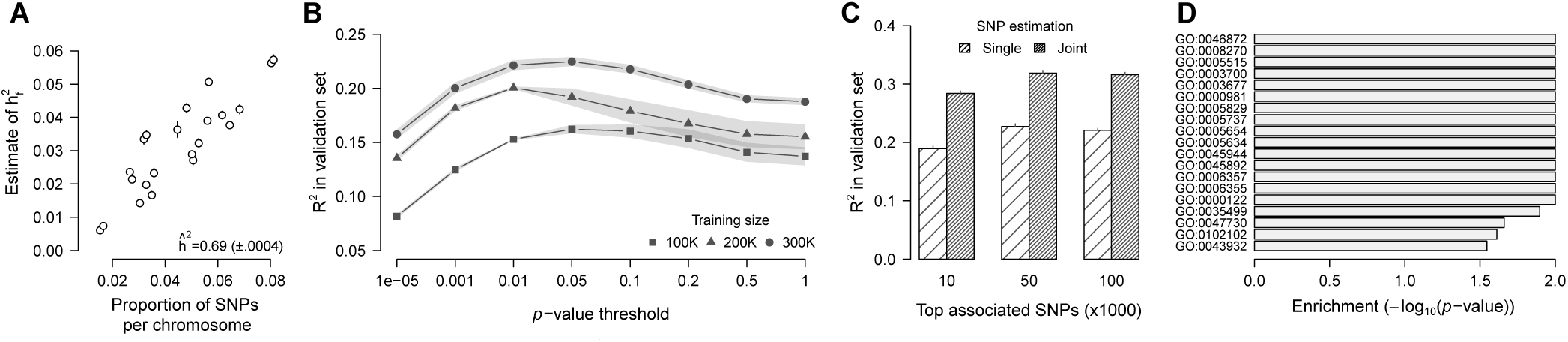
Genetic analysis of human height. (**A**) Estimates of proportion of genetic variance captured by each autosomal chromosome 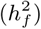 as a function of proportion of SNPs per chromosome. Error bars represent the standard error (SE) obtained from analysing five data subdivisions each containing 50,000 individuals. The heritability (*h*^2^±SE) was estimated to 0.69±0.0004. (**B**) Variance explained (*R*^*2*^) in the validation set by polygenic risk scores (PRS) using a range of single marker SNP *p*-value thresholds (pruned for linkage disequilibrium, *r*^2^ < 0.7). The shaded area depicts the SE of *R*^2^ across the five validation sets. (**C**) Joint estimation of SNP effects of top markers leads to improved *R*^2^ in the validation set compared to PRS using single SNP effects from linear model associations. The re-estimation of top SNPs was performed within training sets of 300K individuals. (**D**) Enrichment of SNPs within genomic features defined by gene ontology (GO) terms. The enrichment analysis was performed using the single marker test statistics from linear model associations on five training sets of 300K individuals; thus, the *p*-values shown here are the average *p*-values across the five training sets, adjusted for multiple testing using a false discovery rate < 0.05.

The qgg package contain a novel approach for computing polygenic risk scores. Using a selected set of top associated SNPs, for example top 10*K*, 50*K* or 100*K*, we re-estimate the SNP effects, but in a model where the SNP effects are jointly estimated (in contrary to traditional linear model association that estimate the effects SNP by SNP). This leads to increased predictive performance from *R*^*2*^= 0.22 to *R*^*2*^= 0.32 (Fig. 1C, see Supplementary Information section 9 for methodological details).

Finally, the SNP effects from single marker associations were used to perform gene set enrichment analysis on genomic features defined by gene ontology (GO) terms. We identified 19 GO terms that had an average false discovery rate < 0.05 across the five training sets (*n* = 300*K*, Fig. 1D); for example the GO term “ossification involved in bone remodelling” (GO:0043932), and several terms related to the synthesis of carnosine (GO:0035499, GO:0047730, GO:0102102); a dipeptide which is naturally occurring in skeletal muscles, and intake of carnosine supplements has been shown to increase the anaerobic performance (Hill *et al*., 2007).

## Conclusion

The qgg package provides an infrastructure for analysing large-scale genotype-phenotype data, and contains a range of quantitative genetic modelling approaches for investigating the genetic basis of complex traits and diseases, including a novel prediction approach which improves the accuracy of prediction compared to commonly used methods.

