## Supplementary information for "qgg: an R package for large-scale quantitative genetic analyses"

---

The qgg package uses a simple infrastructure with a uniform set of arguments across all functions. The general workflow can be divided into two main steps: 1) Prepare data for statistical analyses, and 2) Perform statistical analyses (Figure S1).

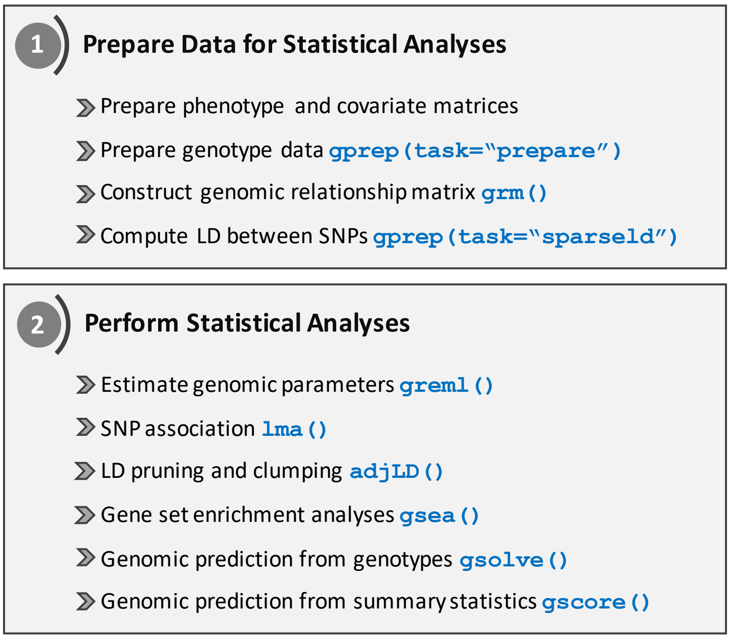

**Figure S1:** Overview of the core facilities that are implemented in the qgg package.

This supplementary file contains details on the methods implemented in the qgg package, and examples on the usage of the main functions. The document is organised into several sections, each describing one of the core facilities in qgg. Each section is concluded with examples. For the examples we use the genetic and phenotypic data (human standing height) from the UK Biobank (UKB) (Bycroft et al., 2018). The Appendix contains the R-code for the examples within the main manuscript.

The qgg package was developed based on the hypothesis that certain regions on the genome, so-called *genomic features*, may be enriched for causal genetic variants affecting the complex trait or disease. Therefore, we begin with an introduction to genomic features, and how these can be established using available resources, such as Bioconductor (Huber et al., 2015).

In the R environment the qgg package can be installed directly from CRAN (which we recommend);

```
install.packages("qgg")
library(qgg)
```

or the latest developer version from Github, but this requires additional tools installed, such as Rtools for Windows (https://cran.r-project.org/bin/windows/Rtools/), or Xcode for Mac OS (https://developer.apple.com/xcode/).

```
install.packages("devtools")
library(devtools)

install_github("psoerensen/qgg")
library(qgg)
```

**10 Computational times** **21**

**Literature cited** **22**

**Appendix: R-code for UKB example** **24**

### 1 Genomic features

#### 1.1 Genomic feature classes

Genomic features are collections of genetic variants (such as single nucleotide polymorphisms (SNPs)) grouped based on common biological- or molecular functions, or other characteristics. The aggregation of genetic variants rely on prior biological knowledge from external sources such as protein-protein interactions (*e.g.* STRING (von Mering et al., 2005)), biological pathways (*e.g.* KEGG (Kanehisa and Goto, 2000)), gene functions (*e.g.* gene ontologies, GO (The Gene Ontology Consortium, 2000)), sequence ontologies (*e.g.* introns, exon and binding sites (Eilbeck et al., 2005)), drug targets (*e.g.* drug bank (Wishart et al., 2006)), genome-wide expression patterns (*e.g.* GTEx (GTEx Consortium, 2015)), or prior trait associations (*e.g.* human GWAS catalog (MacArthur et al., 2017)). In addition, feature sets can be created from other types of omic data such as metabolomic, proteomic or epigenetic variation.

Naturally, only genetic markers located within the genomic features can be considered in the analysis; thus, an important step is the mapping of genetic variants to the genomic features (Figure S2). This is typically done by grouping all markers within known gene regions. To capture the regulatory regions of genes the upstream and downstream regions are often included, and potentially also any regions in linkage disequilibrium (LD) with the gene. Therefore, some markers may be linked to multiple feature sets.

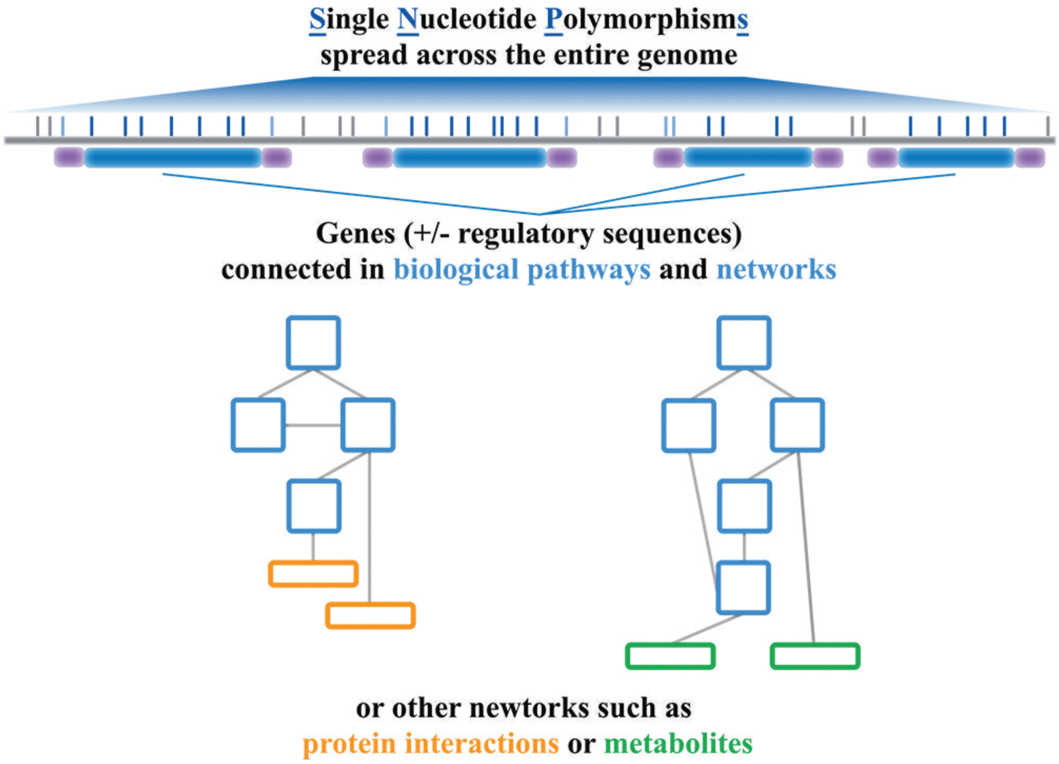

**Figure S2:** Graphical representation of genomic feature classes. First, all SNPs located within the same gene region (*e.g.* the transcribed region, dark-blue SNPs) are aggregated. Gene regions can be extended such that SNPs within regulatory regions are included (light-blue SNPs). Second, genes can then be grouped based on prior biological information, *genomic features*, such as genes connected in pathways or based on other similarities, such as protein networks or common metabolite signatures. The genomic feature classes are thus collection of SNPs that has been aggregated based on shared biological or molecular characteristics.

The degree of new knowledge obtained from the genomic feature analysis strongly depends on the quality and complexity of the genomic feature class. The more reliable the resource is, the more accurate the following results will be. However, if the degree of feature complexity is low, and the quality is high, the outcome might be of minor interests; *e.g.*, chromosomal regions are well-defined features, but enrichment of certain chromosomes or chromosomal regions might be of less interest. It is therefore important prior to the feature analysis to clearly formulate the scope of the analysis.

Genomic feature models have the prospect to contribute with novel knowledge, but they highly depend upon the availability and specificity of the prior biological information. Unfortunately, such information is not readily available across the tree of life. For model organisms and humans, much information is available, but even for well studied organisms, such as livestock species, the amount and level of detail is limited. However, the definitions of genomic features are constantly evolving, and new knowledge will continuously be added, also for those organisms which currently are lacking good feature information.

#### 1.2 Genomic features in R

Here we will demonstrate one approach to annotate SNPs using the UKB data. The first step is to link SNPs to the genes which the SNPs are located within. Such annotation can be done in several ways, but here we will use Biconductor (Huber et al., 2015) and the packages `VariantAnnotation` (Obenchain et al., 2014), `TxDb.Hsapiens.UCSC.hg19.knownGene` and `org.Hs.eg.db`.

```
# install and load packages
  install.packages("data.table")
  install.packages("BiocManager")
  BiocManager::install("VariantAnnotation", version = "3.8")
  BiocManager::install("TxDb.Hsapiens.UCSC.hg19.knownGene", version = "3.8")
  BiocManager::install("org.Hs.eg.db", version = "3.8")

  library(VariantAnnotation); library(TxDb.Hsapiens.UCSC.hg19.knownGene)
  library(org.Hs.eg.db); library(data.table)

# map between Entrez Gene Identifyer and Gene Symbols
  eg2sym <- org.Hs.egSYMBOL
  mapped_genes <- mappedkeys(eg2sym)
  eg2sym <- as.list(eg2sym[mapped_genes])

# read SNPs from PLINK bim-file
  snp <- NULL
  for( chr in 1:22 ){
       fnIn <- fread(paste0("./genotypes/ukb_snp_chr",chr,"_v2.bim"), data.table=FALSE)
       colnames(fnIn) <- c("chr", "rsid", "cm", "bp", "a1", "a2")
       fnIn$chr <- paste("chr", fnIn$chr, sep="")
       snp <- rbind(snp, fnIn)
       print(chr)
  }

# create GRanges object for the SNPs
  assoc <- with(snp, GRanges( seqnames = Rle(chr),
                ranges = IRanges(bp, end=bp, names=rsid), strand = Rle(strand("*"))))

# get entrez gene ids for the SNPs
  loc <- locateVariants(assoc, TxDb.Hsapiens.UCSC.hg19.knownGene,
                        AllVariants(intergenic =  IntergenicVariants(upstream = 5000,
                        downstream = 5000)))

# convert the GRanges object to data frame
  names(loc) <- NULL
  snpA <- as.data.frame(loc)
  snpA$rsid <- names(assoc)[ snpA$QUERYID ]
  snpA <- snpA[, c("rsid", "seqnames", "start", "end", "LOCATION",
                   "GENEID", "PRECEDEID", "FOLLOWID")]
  snpA <- unique(snpA)

# add gene symbols
  sym <- eg2sym[match(snpA$GENEID, names(eg2sym))]
  isNULL <- sapply(sym, is.null)
  sym[isNULL] <- NA
  snpA$sym <- unlist(sym)

  save(snpA, file="./data/snpA.Rdata")

> head(snpA)
           rsid seqnames  start    end   LOCATION GENEID PRECEDEID FOLLOWID  sym
  1  rs28659788     chr1 723307 723307 intergenic   <NA>                    <NA>
  2 rs116587930     chr1 727841 727841 intergenic   <NA>                    <NA>
  3 rs116720794     chr1 729632 729632 intergenic   <NA>                    <NA>
  4   rs3131972     chr1 752721 752721 intergenic   <NA>                    <NA>
  5  rs12184325     chr1 754105 754105 intergenic   <NA>                    <NA>
  6   rs3131962     chr1 756604 756604 intergenic   <NA>     79854          <NA>
```

In the UKB data there are 779,903 SNPs available distributed across 20,775 genes with an average of 24.7 SNPs per gene (min.=1, max.=2,117, Figure S3).

The genes can now be grouped based on shared characteristics, such as KEGG pathways (Kanehisa and Goto, 2000) or gene ontology categories (The Gene Ontology Consortium, 2000). For this, we can use the Bioconductor package `org.Hs.eg.db`.

```
### SNPs --> Gene ontologies
  go2eg <- org.Hs.egGO2EG
```

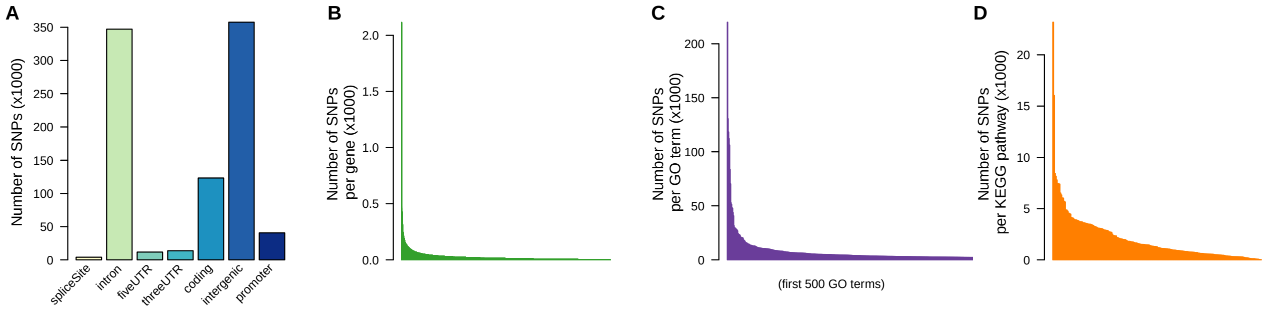

**Figure S3:** Annotation of UKB SNPs on autosomal chromosomes. **A:** Distribution of SNPs in different sequence ontology classes. **B:** Number of SNPs across the 20,775 genes. **C:** Number of SNPs linked to gene ontology (GO) terms (only the 500 GO terms with most SNPs are shown), and **D:** number of SNPs per KEGG pathway.

```
  mapped_genes <- mappedkeys(go2eg)
  go2eg <- as.list(go2eg[mapped_genes])

   goSets <- lapply(go2eg, function(x){
      eg <- x
      eg <- eg[eg%in%snpA$GENEID]
      snp <- unique(snpA[snpA$GENE%in%eg,]$rsid )
      return(snp)
  })

# remove units with less than five SNPs
  isNULL <- sapply(goSets, is.null)
  goSets <- goSets[!isNULL]
  l <- sapply(goSets, length)
  goSets <- goSets[l>4]
  save(goSets, file="./data/goSets.Rdata")

> str(goSets)
  List of 6218
  $ GO:0000002: chr [1:67] "rs12140494" "rs72646720" "rs77189728" "rs115161668" ...
  $ GO:0000003: chr [1:12] "rs2377027" "rs138510303" "Affx-52323116" "rs2889537" ...
  $ GO:0000027: chr [1:40] "rs1171" "Affx-80210241" "rs11548045" "rs2275915" ...
  $ GO:0000028: chr [1:5] "rs188748675" "rs41265211" "rs41265213" "rs114120114" ...
  $ GO:0000045: chr [1:69] "rs112485554" "rs72655053" "rs72655055" "rs114855850" ...
  $ GO:0000052: chr [1:66] "rs233114" "rs1498374" "Affx-80267987" "Affx-80212595" ...

### SNPs --> KEGG pathways
  kegg2eg <- org.Hs.egPATH2EG
  mapped_genes <- mappedkeys(kegg2eg)
  kegg2eg <- as.list(kegg2eg[mapped_genes])

  keggSets <- lapply(kegg2eg, function(x){
      eg <- x
      eg <- eg[eg%in%snpA$GENEID]
      snp <- unique(snpA[snpA$GENE%in%eg,]$rsid )
      return(snp)
  })

# remove units with less than five SNPs
  isNULL <- sapply(keggSets, is.null)
  keggSets <- keggSets[!isNULL]
  l <- sapply(keggSets, length)
  keggSets <- keggSets[l>4]
  save(keggSets, file="./data/keggSets.Rdata")

> str(keggSets)
  List of 208
  $ 04610: chr [1:395] "Affx-92047482" "Affx-80267243" "rs137990115" "Affx-92047483" ...
  $ 00983: chr [1:252] "rs603412" "rs2072671" "rs818202" "rs12143372" ...
  $ 01100: chr [1:2545] "rs7796" "Affx-89008512" "rs149269930" "Affx-5751253" ...
  $ 00380: chr [1:88] "rs4233335" "rs144984854" "rs77951676" "rs200550194" ...
  $ 00970: chr [1:109] "Affx-80267535" "rs141323599" "Affx-80211368" "rs10914614" ...
```

These sets can then for example be used in the gene set enrichment analyses (see Section 6)

### 2 Prepare phenotype data

#### 2.1 Streamline data

The preparation of the phenotypic data will be unique for each study, but a general recommendation is to prepare and streamline the phenotype and covariate data to avoid manipulation of the data for each analysis. This will likely reduce the chance of errors, and make the scripts as simple as possible. We also recommend to prepare training and validation sets at this point.

#### 2.2 Phenotype data from UKB

The phenotypic data from UKB can be manipulated with the R package `ukbtools` (Hanscombe et al., 2017). After the UKB data have been downloaded and decrypted with the supplied UKB programs (http://biobank.ndph.ox.ac.uk/showcase/download.cgi), the UKB dataset can be constructed with the function `ukb_df` from the `ukbtools` package:

```
# install and load packages
  install.packages("ukbtools")
  library(ukbtools)
  ukb_data <- ukb_df("ukbxxxx", path="./ukb_rawfiles/")

  save(ukb_data, file="./phenotypes/ukb_data.Rdata")
```

Next, we restrict the phenotype file (`ukb_data`) to the White British cohort of unrelated individuals with no aneuploidy and $\leq 5\%$ missing genotypes.

```
  load(file="./phenotypes/ukb_data.Rdata")
  df <- ukb_data

# relatedness information
  rel <- read.table(file="./genotypes/ukbxxxxxx_rel_syyyyyy.dat",sep=" ", header=TRUE)

# extract the White British cohort
  is.British <- df[, "ethnic_background_0_0" ]=="British"
  df <- df[is.British,]
  is.white <- df[,"genetic_ethnic_grouping_0_0"]=="Caucasian"
  is.white[is.na(is.white)] <- FALSE
  df <- df[is.white,]

# remove related individuals
  ids1 <- rel[,"ID1"]
  ids2 <- rel[,"ID2"]
  ids_related <- ids2[!ids2%in%ids1]
  ids_related <- ids2

  is.related <- df[,"eid"]%in%ids_related
  df <- df[!is.related,]

# remove individuals with aneuploidy
  is.aneuploidy <- df[,"sex_chromosome_aneuploidy_0_0"]=="Yes"
  is.aneuploidy[is.na(is.aneuploidy)] <- FALSE
  df <- df[!is.aneuploidy,]

# remove individuals with genotype missingness above 0.05
  is.missing <- df[,"missingness_0_0"]<=0.05
  df <- df[is.missing,]

  df$eid <- as.character(df$eid)
  rownames(df) <- df$eid
```

The data frame (`df`) contains many more variables than what is needed here; thus, we generate a data frame that only contains the phenotype (standing height) and selected covariates (sex, age, and the ten first genetic principal components).

```
  cls <- c("sex_0_0",
           "year_of_birth_0_0",
           "genetic_principal_components_0_1",
           "genetic_principal_components_0_2",
           "genetic_principal_components_0_3",
           "genetic_principal_components_0_4",
           "genetic_principal_components_0_5",
           "genetic_principal_components_0_6",
```

```
            "genetic_principal_components_0_7",
            "genetic_principal_components_0_8",
            "genetic_principal_components_0_9",
            "genetic_principal_components_0_10",
            "standing_height_0_0")

  df <- df[,cls]
  colnames(df) <- c("sex","age",paste("pc",1:10,sep=""), "height")
  df[,"age"] <- 2018-df[,"age"]

# centre and scale the phenotype (mean=0, sd=1)
  df$height <- scale(df$height)

  save(df,file="./phenotypes/df_wbu_ukb.Rdata")
```

The phenotype is then corrected for the covariates, which we do using the regression function `fastlm` that is implemented in the qgg package.

```
# adjust y for covariates
  df$height_scaled <- NA
  isNA <- is.na(df[,"height"])
  X <- model.matrix(~ sex + age + pc1 + pc2 + pc3 + pc4 + pc5 + pc6 + pc7 + pc8 + pc9 + pc10,
                    data=df[!isNA,])
  fit <- fastlm(y=df[!isNA,"height"],X=X)
  df[!isNA,"height_scaled"] <- df[!isNA,"height"]-fit$yhat

  save(df,file="./phenotypes/df_wbu_ukb_scaled.Rdata")
```

The last step in phenotype preparation is to establish training and validation sets. For illustration purposes, we here use a five-fold cross validation scheme consisting of a total of 25,000 individuals. The `R` package `caret` (Kuhn, 2008) provide a simple solution to this (see Appendix for other alternatives)

```
# install and load packages
  install.packages("caret")
  require(caret)

# define CV-scheme
  N <- 25000
  df <- df[!is.na(df$height),]
  df <- df[sample(1:nrow(df), N),]
  ids <- rownames(df)
  train <- createFolds(ids, k=5, returnTrain=TRUE)
  train <- lapply(train, function(x){ ids[x] })

  ytrain <- NULL
  for( i in 1:length(train) ){
       yobs <- df$height_scaled
       names(yobs) <- rownames(df)
       isT <- names(yobs)%in%train[[i]]
       yobs[!isT] <- NA
       ytrain <- cbind(ytrain,as.matrix(yobs))
       print(i)
  }
  colnames(ytrain) <- names(train)

  save(ytrain,file="./phenotypes/ytrain.Rdata")
  save(train,file="./phhenotypes/train.Rdata")
```

### 3 Prepare genotype data

#### 3.1 Large-scale genetic data

The final step in data preparation is to prepare a genotype file. For large-scale genetic data it can be challenging to work with the entire data set loaded in memory. One solution is to process the genotypes directly. Many genetic data sets are stored in the binary PLINK format (Purcell et al., 2007). The qgg package provides functions that work directly with binary PLINK files. If the genetic data can be in memory (*i.e.*, small data sets) it may be more convenient to work directly on the loaded files. In such cases the small data set tutorial may be helpful (see `Small data set tutorial` on our webpage).

##### 3.2 Processing genotype data with qgg

To prepare the genotype file (for large-scale genetic data) for downstream analysis, the binary PLINK file(s) has to be processed. This is done with the function `gprep`. This function has two purposes; 1) convert the PLINK file(s) to a `*.raw`-format, and 2) creates an `R`-object that contains information about the current study, such as individual and SNP IDs, path to PLINK- and `*.raw`-files, and summary information like minor allele frequencies (MAF). If the original binary data file contains more individuals and SNPs than needed for the downstream analyses, this can be specified with the arguments `ids=` and `rsids=`. The function `gprep` creates a new genotype file (`*.raw`, path specified with argument `fnRAW=`) that contains the genotypes as allele counts of the alternative allele (memory usage = $(n \text{ x } m)/4$ bytes).

```
# load individual IDs
  load(file="./phenotypes/df_wbu_ukb.Rdata")
  ids <- rownames(df)

# path to PLINK file
  bimfiles <- paste("./genotypes/ukb_snp_chr",1:22,"_v2.bim",sep="")
  bedfiles <- paste("./genotypes/ukb_cal_chr",1:22,"_v2.bed",sep="")
  famfiles <- paste("./genotypes/ukbxxxxxx_cal_chr",1:22,"_v2_szzzzzz.fam",sep="")

# path to raw-file
  fnRAW <- "./genotypes/wbu_ukb.raw"

# prepare the genotype file
  Glist <- gprep(study="WBU_UKB", fnRAW=fnRAW, bedfiles=bedfiles, bimfiles=bimfiles,
                famfiles=famfiles, ids=ids)
  save(Glist, file="./genotypes/Glist_wbu_ukb.Rdata", compress=FALSE)
```

The object `Glist` contains all the relevant information about the current study; including minor allele frequencies (`maf`), number of individuals with missing genotypes per genotype (`nmiss`), homozygozity (`hom`) and heterozygosity (`het`), and the number of the genotype classes per loci (`n0`, `n1`, `n2`).

```
> names(Glist)
   [1] "fnRAW"       "rsids"       "alleles"     "position"    "af"
   [6] "maf"         "nmiss"       "het"         "n0"          "n1"
  [11] "n2"          "chr"         "chrnames"    "nchr"        "ids"
  [16] "study_ids"   "study_rsids" "mchr"        "m"           "n"
  [21] "study"       "bedfiles"    "bimfiles"    "famfiles"    "hom"
```

The slot `Glist$study_rsids` can be used to store the SNP IDs of those genotype markers that should be used in the downstream analyses; for example, those that have passed filtering criteria such as minor allele frequency $\geq 0.01$, the proportion of missing genotypes $< 5\%$ and exclusion of particular genomic regions, such as the major histocompatibility complex:

```
  isMAF <- Glist$maf<=0.01
  isMISS <- Glist$nmiss/length(Glist$study_ids) > 0.05
  isMHC <- Glist$chr==6 & Glist$position > 25602429 & Glist$position < 33471466
  delete <- isMAF | isMISS | isMHC

  Glist$study_rsids <- Glist$rsids[!delete]
```

A subset of SNP genotypes can be read into `R` with the function `getW`.

```
  W <- getW(Glist=Glist, ids=Glist$ids[1:5], rsids=Glist$study_rsids[1:6])
> W
      rs28659788 rs116587930 rs116720794 rs3131972 rs12184325 rs3131962
  id1          0           0           0         2          0         2
  id2          0           0           0         2          0         2
  id3          0           1           0         1          0         1
  id4          0           0           0         2          0         2
  id5          0           0           0         2          0         2
```

For the UKB example we have a total of $n = 335,744$ individuals and $m = 599,297$ SNPs for downstream analyses.

#### 4 Estimation of genetic parameters

##### 4.1 Variance component estimation

Estimation of heritability ($\hat{h}^2$) of complex traits and diseases can be achieved by estimating the variance components with Restricted Maximum Likelihood (REML) in the Genomic Best Linear Unbiased Prediction (GBLUP) framework.

$$\mathbf{y} = \mathbf{X}\mathbf{b} + \mathbf{Z}\mathbf{g} + \mathbf{e} \text{ with } \mathrm{Var}(\mathbf{y}) = \mathbf{G}\sigma_g^2 + \mathbf{I}\sigma_e^2, \tag{1}$$

where $\mathbf{y}$ is an $n \times 1$ vector of phenotypic values with $n$ being the sample size, $\mathbf{b}$ is a vector of fixed effects (like sex, age, and genetic principal components), $\mathbf{g}$ and $\mathbf{e}$ are $n \times 1$ vectors of total genetic and residual effects, respectively; both assumed to be independent normally distributed as: $\mathbf{g} \sim N(0, \mathbf{G}\sigma_g^2)$ and $\mathbf{e} \sim N(0, \mathbf{I}\sigma_e^2)$. The genetic relationship between the individuals is captured by the additive genomic relationship matrix (GRM, $\mathbf{G}$), which can be constructed from the SNP markers (VanRaden, 2008; Yang et al., 2011): $\mathbf{W}\mathbf{W}'/m$, where $\mathbf{W}$ is the centred and scaled genotype matrix, and $m$ is the number of markers. Each column vector of $\mathbf{W}$ is: $\mathbf{w}_i = (\mathbf{m}_i - 2p_i)/\sqrt{2p_i(1-p_i)}$, where $p_i$ is the minor allele frequency of the $i$th genetic marker, and $\mathbf{m}_i$ is the $i$th column vector of the allele count matrix, $\mathbf{M}$, which contains the genotype coded as $\{0, 1, 2\}$ counting the number of minor allele.

Two other types of genomic relationship matrices can be defined; the epistatic genomic relationship matrix ($\mathbf{G}_{aa}$) and the dominance genomic relationship matrix ($\mathbf{D}$). The epistatic genomic relationship matrix can be derived from the additive genomic relationship matrix (when considering first order epistatic interactions, *i.e.* additive by additive) as: $\mathbf{G}_{aa} = \mathbf{G}\#\mathbf{G}$ (Su et al., 2012), where # denotes the Hadamard product operation. The dominance genomic relationship matrix can be obtained as: $\mathbf{D} = \mathbf{H}\mathbf{H}'/\sum 2p_iq_i(1-2p_iq_i)$, where $\mathbf{H}$ is a $n \times m$ matrix of heterozygosity coefficients with element $h_{k,i} = 0 - 2p_iq_i$ if individual $k$ is homozygous, and $h_{k,i} = 1 - 2p_iq_i$ if individual $k$ is heterozygous at locus $i$ (Su et al., 2012).

The variance components, *i.e.* $\theta = (\sigma_{g_1}^2, \sigma_{g_2}^2, \ldots, \sigma_{g_k}^2, \sigma_e^2)$, can be estimated using restricted maximum likelihood (REML). The restricted log-likelihood corresponding to the model in Equation 1 can be written as:

$$\mathbf{L} = -0.5(\log|\mathbf{V}|) + \log(|\mathbf{X}'\mathbf{V}\mathbf{X}|) + \mathbf{y}'\mathbf{P}\mathbf{y}, \tag{2}$$

where $\mathbf{P} = \mathbf{V}^{-1} - \mathbf{V}^{-1}\mathbf{X}(\mathbf{X}'\mathbf{V}^{-1}\mathbf{X})^{-1}\mathbf{X}'\mathbf{V}^{-1}$ is the projection matrix projecting the observed phenotypes into residuals, $\mathbf{e} = \mathbf{P}\mathbf{y}$. Also note that $\mathbf{P}\mathbf{y} = \mathbf{V}^{-1}(\mathbf{y} - \mathbf{X}\hat{\mathbf{b}})$, where $\hat{\mathbf{b}} = (\mathbf{X}'\mathbf{V}^{-1}\mathbf{X})^{-1}\mathbf{X}'\mathbf{V}^{-1}\mathbf{y}$.

The variance components, $\theta$, can be estimated using the average information (AI) REML algorithm, which is based on an iterative procedure:

$$\theta^{j+1} = \theta^j + (\mathbf{AI})^{-1}\mathbf{s}, \tag{3}$$

where $\theta^j$ is the vector of estimated variance components in iteration $j$, and $\mathbf{s}$ is the vector of first derivatives of the restricted log-likelihood with respect to the variance component,

$$\mathbf{s}_i = \frac{1}{2}\left[tr\Big(\mathbf{P}(\frac{\partial \mathbf{V}}{\partial \sigma_i^2})\Big) - \mathbf{y}'\mathbf{P}(\frac{\partial \mathbf{V}}{\partial \sigma_i^2})\mathbf{P}\mathbf{y}\right]. \tag{4}$$

$\mathbf{AI}$ is called the average information matrix, which is the average of the observed and expected information matrix based on second derivatives of the restricted log-likelihood, where the $ij$-elements are obtained as,

$$\mathbf{AI}_{ij} = \frac{1}{2}\Big(\mathbf{y}'\mathbf{P}\frac{\partial \mathbf{V}}{\partial \sigma_i^2}\mathbf{P}\frac{\partial \mathbf{V}}{\partial \sigma_j^2}\mathbf{P}\mathbf{y}\Big). \tag{5}$$

The iterative process is terminated when the change in updating of parameters are sufficiently small, *e.g.* $||\theta^j - \theta^{j+1}|| < \epsilon$, where $\epsilon$ is a very small value, for example $1 \times 10^{-8}$.

The model in Equation 1 can be extended to include multiple genetic components; for example, to estimate the proportion of explained genetic variance by SNPs located in a specific genomic feature or across several features, like autosomal chromosomes.

$$\mathbf{y} = \mathbf{X}\mathbf{b} + \sum_{k=1}^{n_f} \mathbf{Z}\mathbf{g}_k + \mathbf{e}, \tag{6}$$

where $\mathbf{g}_k$ is the genetic values captured by the $k$-th marker set with the additive genomic relationship matrix computed for the subset of markers as $\mathbf{G}_k = \frac{\mathbf{W}_k\mathbf{W}_k'}{m_k}$. The phenotypic variance then becomes: $\mathbf{V} = \sum_{k=1}^{n_f} \mathbf{Z}\mathbf{G}_k\mathbf{Z}'\sigma_{g_k}^2 + \mathbf{I}\sigma_e^2$.

The proportion of phenotypic variance explained by additive genetic effects is the narrow sense heritability (Falconer and Mackay, 1996; Lynch and Walsh, 1998),

$$h^2 = \frac{\sigma_a^2}{\sigma_p^2}, \tag{7}$$

where $\sigma_a^2$ is the additive genetic variance, and $\sigma_p^2$ is the total phenotypic variance. If genome-wide SNP data is applied one can estimate the proportion of phenotypic variance captured by the SNPs (Yang et al., 2010; de los Campos et al., 2015)

$$\hat{h}_g^2 = \frac{\hat{\sigma}_g^2}{\hat{\sigma}_p^2}. \tag{8}$$

For models with multiple random genetic effects, the genomic heritability can be expressed as,

$$\hat{h}_g^2 = \frac{\sum_{i=1}^{n_f} \hat{\sigma}_{g_1}^2 + \hat{\sigma}_{g_2}^2 + \cdots + \hat{\sigma}_{g_{n_f}}^2}{\sum_{i=1}^{n_f} \hat{\sigma}_{g_1}^2 + \hat{\sigma}_{g_2}^2 + \cdots + \hat{\sigma}_{g_{n_f}}^2 + \hat{\sigma}_e^2}. \tag{9}$$

In the case of a two component feature model, such as $\mathbf{y} = \mathbf{f} + \mathbf{r} + \mathbf{e}$ (Eq. 6), where $\mathbf{f}$ and $\mathbf{r}$ indicates the random genetic effects captured by the SNPs within and outside the genomic feature, respectively, the genomic heritability is,

$$\hat{h}_g^2 = \frac{\hat{\sigma}_f^2 + \hat{\sigma}_r^2}{\hat{\sigma}_f^2 + \hat{\sigma}_r^2 + \hat{\sigma}_e^2}. \tag{10}$$

Then, the proportion of total genetic variance captured ($\hat{h}_f^2$) by the genetic markers in the genomic feature can be expressed as,

$$\hat{h}_f^2 = \frac{\hat{\sigma}_f^2}{\hat{\sigma}_f^2 + \hat{\sigma}_r^2}. \tag{11}$$

#### 4.2 Genomic prediction

Beside estimating the genetic contribution to complex traits, we can also predict the genetic values given the estimated parameters. If, the data set is split into a training ($t$) and validation ($v$) set, we estimate the variance components using the training population, and use those parameters and the co-variance among the individuals (given by the GRM) to predict the genetic values for the individuals in the validation population:

$$\hat{\mathbf{g}}^v = (\sum_{i=1}^{n_f} \mathbf{G}_i^{vt} \sigma_{g_i}^2)[\sum_{i=1}^{n_f} \mathbf{G}_i^{tt} \sigma_{g_i}^2 + \mathbf{I}^{tt} \sigma_e^2]^{-1} (\mathbf{y}^t - \mathbf{X}^t \hat{\mathbf{b}}^t), \tag{12}$$

where $i$ is the i-$th$ genetic components ($n_f$), and the additive genomic relationship matrices,

$$\mathbf{G}_i = \begin{pmatrix} \mathbf{G}_i^{vv} & \mathbf{G}_i^{vt} \\ \mathbf{G}_i^{tv} & \mathbf{G}_i^{tt} \end{pmatrix}, \tag{13}$$

are partitioned according to relationships between the individuals in the training set ($\mathbf{G}_i^{tt}$), between the individuals in the validation set ($\mathbf{G}_i^{vv}$), and among individuals in the training and validation set ($\mathbf{G}_i^{vt}$).

The predictive performance can then be summarised as the correlation between the observed phenotype and the predicted genetic values, $Cor(\mathbf{y}^v, \hat{\mathbf{g}}^v)$, or as the proportion of variance explained in the validation set by a linear fit ($R^2$). For binary traits (such as diseases) the accuracy of prediction can be quantified using Nagelkerke's $R^2$,

$$R^2 = \frac{1 - e^{(-LR/n)}}{1 - e^{-(-2L_0)/n}}, \tag{14}$$

where $n$ is the number of observations, $LR$ is the likelihood ratio comparing two nested logistic regression models, and $L_0$ is the log-likelihood of the null model neglecting the predicted genetic effects as predictor. Alternatively, we can use the area under the ROC Curve (AUC) to assess the degree of correct classification of cases and controls (Wray et al., 2010). According to Wray et al. (2010), AUC can be computed as

$$\text{AUC} = \frac{1}{n_{d'}}(\bar{r}_d - \frac{n_d}{2} - \frac{1}{2}), \tag{15}$$

where $\bar{r}_d$ is the mean rank of cases, and $n_d$ and $n_{d'}$ are the number of cases and controls, respectively. Thus, after estimating the genetic risk, the individuals are ranked according to their disease risk (the individual with the highest predicted risk has rank $r_1 = n_d + n_{d'} = n$). AUC lies between zero and one, where one is the upper bound indicating perfect ranking of cases and controls.

#### 4.3 Constructing GRM with `grm`

In the qgg package the additive genomic relationship matrix is constructed with the function `grm`. Similar to the `gprep` function, `grm` creates two items, the GRM on disk, and an R-object `GRMlist`, which contains all essential information about that specific GRM. If the GRM should be centred and scaled (*i.e.*, each column vector of `W` has a mean of zero and variance of one) use the argument `scale=TRUE`.

```
# compute GRM for the individuals in training set 1 using all SNPs
  idsG <- train[[1]]
  rsids <- Glist$study_rsids

# path to the *.grm-file
  fnG <- "./grm/Gcv1.grm"
  GRMlist <- grm( Glist=Glist, ids=idsG, rsids=rsids, scale=TRUE, msize=100, ncores=12, fnG=fnG)
  save(GRMlist, file="./grm/GRMlistCV1.grm")
```

To view parts of the GRM, we can read the GRM into `R` with the `getGRM` function:

```
  G <- getGRM(GRMlist, ids=ids[1:1000])

# the dimension of the GRM
> dim(G)
  [1] 1000 1000

# diagonal elements
> head(diag(G))
          1         2         3         4         5         6
  0.9895776 1.0105822 1.0109172 1.0105491 0.9926939 0.9936836

# GRM of first five individuals
> G[1:5,1:5]
                1             2             3             4             5
  1  0.9895775614 -0.0031241397  0.0003626016 -0.0008579677  0.0019926350
  2 -0.0031241397  1.0105821983 -0.0029835079 -0.0005884170  0.0005135084
  3  0.0003626016 -0.0029835079  1.0109171969  0.0045359541 -0.0034589046
  4 -0.0008579677 -0.0005884170  0.0045359541  1.0105491427  0.0015048236
  5  0.0019926350  0.0005135084 -0.0034589046  0.0015048236  0.9926938747
```

To compute the non-additive genomic relationship matrices, use the argument `method="dom"` or `method="epi-hadamar"` for the dominance and epistatic genomic relationship matrices, respectively.

For estimation of genetic variance of multiple random genetic effects several GRMs has to be constructed, *e.g.*, for estimating the proportion of genetic variance captured by bins of minor allele frequencies. First, the SNPs are grouped by their MAF, then we construct one GRM for each set of SNPs, and lastly, we construct one `GRMlist` that contains the information for all the GRMs.

```
# divide SNPs by MAF
  rsids <- Glist$study_rsids

  rs1 <- names(Glist$maf[Glist$maf>=0.01 & Glist$maf<=0.05])
  rs1 <- rs1[rs1%in%rsids]

  rs2 <- names(Glist$maf[Glist$maf>0.05 & Glist$maf<=0.1])
  rs2 <- rs2[rs2%in%rsids]

  rs3 <- names(Glist$maf[Glist$maf>0.1 & Glist$maf<=0.2])
  rs3 <- rs3[rs3%in%rsids]

  rs4 <- names(Glist$maf[Glist$maf>0.2 & Glist$maf<=0.3])
  rs4 <- rs4[rs4%in%rsids]

  rs5 <- names(Glist$maf[Glist$maf>0.3 & Glist$maf<=0.4])
  rs5 <- rs5[rs5%in%rsids]

  rs6 <- names(Glist$maf[Glist$maf>0.4 & Glist$maf<=0.5])
  rs6 <- rs6[rs6%in%rsids]

  rs <- list(rs1=rs1, rs2=rs2, rs3=rs3, rs4=rs4, rs5=rs5, rs6=rs6)

# construct one GRM for each set of SNPs
  for( i in 1:length(rs) ){
       fnG <- paste0("./grm/Gmaf",i,".grm")
       GRMlist <- grm(Glist=Glist, ids=idsG, rsids=rs[[i]], scale=TRUE, msize=100,
                      ncores=12, fnG=fnG, overwrite=TRUE)
       save(GRMlist, file=paste0("./grm/GRMlistMAF",i,".Rdata"), compress=FALSE)
  }

# merge the GRMlists to one GRMlist
  GRMlistMAF <- NULL
  for( i in 1:6 ){
       load(paste0("./grm/GRMlistMAF",i,".Rdata"))
```

```
    GRMlist$fnG <- paste0("./grm/Gmaf",i, ".grm")
    GRMlistMAF[[i]] <- GRMlist
}
names(GRMlistMAF)<- paste0("MAF",1:6)
GRMlistMAF <- mergeGRM( GRMlistMAF )
save(GRMlistMAF, file="./grm/GRMlistMAF.Rdata", compress=FALSE)
```

#### 4.4 Estimation with `greml`

With the GRMs constructed we can use the function `greml` to estimate the variance components.

```
# load data
load(file="./grm/GRMlistCV1.grm")
load(file="./phenotypes/df_wbu_ukb.Rdata")

# define input files
idsG <- train[[1]]
y <- df[idsG, "height"]
X <- model.matrix(~ sex + age + pc1 + pc2 + pc3 + pc4 + pc5 + pc6 + pc7 + pc8 + pc9 + pc10,
                  data=df[idsG,])

# run greml
fit <- greml(y=y, X=X, GRMlist=GRMlist, theta=c(0.25,0.05), ncores=12, interface="fortran",
             maxit=10, verbose=TRUE)
```

The estimated variance components are stored in the slot `$theta`.

```
> fit$theta
      G1        E
28.53068 11.22306
```

Thus, using a subset of 20,000 individuals the estimated heritability for human standing height is $\hat{h}^2 = \frac{\hat{\sigma}_g^2}{\hat{\sigma}_g^2+\hat{\sigma}_e^2} = \frac{28.53}{28.53+11.22} = 0.72$.

For models with multiple random genetic effects simply use the merged `GRMlist`:

```
load(file="./grm/GRMlistMAF.Rdata")
fit <- greml(y=y, X=X, GRMlist=GRMlistMAF, theta=rep(0.05,7), ncores=12,
             interface="fortran", maxit=10, verbose=TRUE)
```

Figure S4 shows the proportion of genetic variance captured by each MAF bin (expressed pr SNP).

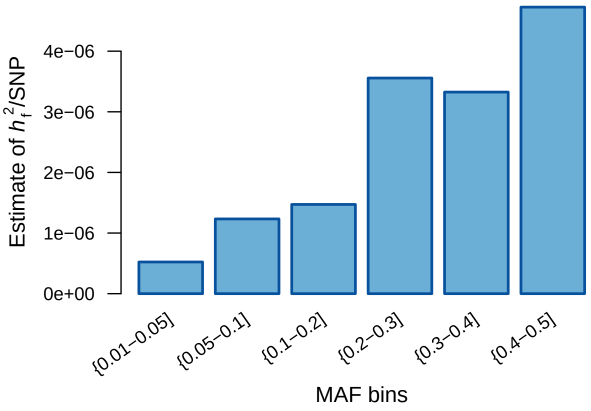

**Figure S4:** Estimates of proportion of genetic variance captured by each MAF bin adjusted for the number of SNPs within each bin.

For small data sets, it is possible to run the analysis within `R` by using the argument `interface="R"`; see tutorial on webpage; Prediction using GBLUP and cross validation.

##### 4.5 Cross-validation example

To illustrate prediction with GBLUP, we use the five fold cross-validation scheme prepared under *"Prepare phenotype data"*. The workflow is, 1) construct the GRM for all individuals (`grm`), 2) estimate variance components for the five different data subdivisions (`greml`), 3) predict the genetic values of the individuals within the validation set (`gblup`), and 4) obtain measures of the predictive performance (`accuracy`).

```
# 1: construct GRM for all 25K
  fnG <- "./grm/G25K.grm"
  idsG <- rownames(ytrain)
  rsids <- Glist$study_rsids
  GRMlist <- grm(Glist=Glist, ids=idsG, rsids=rsids, scale=TRUE, msize=100,
                 ncores=12, fnG=fnG, overwrite=TRUE)
  save(GRMlist, file="./grm/GRMlist25K.grm")

# 2: estimate variance components for the five training sets
  load(file="./phenotypes/df_wbu_ukb.Rdata")
  fit <- NULL
  for( i in 1:length(train) ){
       idsT <- train[[i]]
       y <- df[idsT, "height"]
       names(y) <- rownames(df[idsT,])
       X <- model.matrix(~ sex + age + pc1 + pc2 + pc3 + pc4 + pc5 + pc6 + pc7 + pc8 + pc9 +
                         pc10, data=df[idsT,])
       fit[[i]] <- greml(y=y, X=X, GRMlist=GRMlist, theta=c(0.25,0.05),
                         ncores=12, interface="fortran", maxit=10, verbose=TRUE)
 }

# 3: predict the genetic values for the individuals in the validation sets
  ghat <- NULL
  ids <- GRMlist$idsG
  for( i in 1:length(fit) ){
       idsT <- train[[i]]
       idsV <- ids[!ids%in%idsT]
       ghat[[i]] <- gblup(GRMlist=GRMlist, fit=fit[[i]], ids=idsV)
  }

# 4: accuracy of prediction
  accuracy <- NULL
  for( i in 1:length(ghat) ){
       ypred <- ghat[[i]][,1]
       yobs <- df[names(ypred),"height_scaled"]
       accuracy <- rbind(accuracy,acc(yobs=yobs,ypred=ypred))
  }
```

The mean predictive performance (*i.e.*, $Cor(\mathbf{y}^v, \hat{\mathbf{g}}^v)$) was 0.294 with an average variance explained ($R^2$) in the validation set of $R^2 = 0.079$.

#### 5 Single marker analysis

##### 5.1 SNP-by-SNP association

One of the fundamental approaches to investigate the genetic basis of any complex trait or disease is the single marker analysis, or more commonly known as genome-wide association study (GWAS). This class of methods are hypothesis-free and are used to identify associations between genetic variants and trait variation. Several variations exist, but two main categories are recognised; the linear model analysis (LMA) and the mixed linear model analysis (MLMA). Many different software exists for performing single SNP association analyses, but PLINK (Purcell et al., 2007) is probably the most commonly used.

##### 5.2 SNP association with qgg

Currently there are two approximations for linear and linear mixed model SNP associations implemented in qgg; these are called with the function `lma`. Below we also provide a utility function that allow users to call PLINK (require PLINK to be installed, Purcell et al. (2007)) and run the single SNP analysis.

The PLINK utility function is (not implemented in qgg; thus, the function below has to be copied into the R session). If the trait is binary (*e.g.* disease status), use the argument `logistic=TRUE`.

```
plink_reg <- function(bedfile=NULL, bimfile=NULL, famfile=NULL, yFile=NULL, covFile=NULL,
                      covar=NULL,fids=NULL, ids=NULL, rsids=NULL, fnSNPs=NULL, fnOut=NULL,
                      wd=NULL, logistic=FALSE){
    #bedfile: path to bedfile
    #bimfile: path to bimfile
    #famfile: path to famfile
    #yFile: path to phenotype file
    #fids: family IDs
    #id: ID
    #rsids: vector of SNPs used in GWAS
    #fnSNPs: if you already have a list stored on disk
    #covFile: matrix containing fids, ids, and covariates used in GWAS
    #covar: list which covariables used in the analysis - should be in one character string; fx "
        sex pc1 pc2 pc3", and not "sex", "pc1", "pc2" "pc3"
    #fnOut: path+name of GWAS results
    #wd: working directory where the job should put temporary files
    #logistic: whether the regression should use the link function for logistic regression

  fnY <- paste0(wd,"y")
  fnIDs <- paste0(wd, "id")
  fnCOV <- paste0(wd,"cov")

  write.table( yFile, row.names = F, col.names = T, quote = F, sep = "\t", file = fnY  )
  write.table( cbind(fids,ids), row.names = F, col.names = F, quote = F, sep = "\t", file = fnIDs
        )
  write.table( covFile, row.names = F, col.names = T, quote = F, sep = "\t", file = fnCOV )

  if(!is.null(rsids)){
      fnSNPs <- paste0(wd, "rsids")
      write.table( rsids, row.names = F, col.names = F, quote = F, sep = "\t", file = fnSNPs )
  }

  if(logistic==TRUE){
    system( paste("plink --bfile", bedfile,
                  "--fam", famfile,
                  "--bim", bimfile,
                  "--keep", fnIDs,
                  "--extract", fnSNPs,
                  "--logistic hide-covar",
                  "--keep-allele-order",
                  "--covar", fnCOV,
                  "--covar-name", covar,
                  "--pheno", fnY,
                  "--silent",
                  "--out", fnOut ) )
    }

  if(logistic==FALSE){
    system( paste("plink --bfile", bedfile,
                  "--fam", famfile,
                  "--bim", bimfile,
                  "--keep", fnIDs,
                  "--extract", fnSNPs,
                  "--linear hide-covar",
                  "--keep-allele-order",
                  "--covar", fnCOV,
                  "--covar-name", covar,
                  "--pheno", fnY,
                  "--silent",
                  "--out", fnOut ) )
    }
}
```

Below is an example where we ran the GWAS using the PLINK utility function on $300K$ individuals in the UKB data.

```
  wd <- "./results/"

# get 300K individuals
  df$FID <- df$IID <- rownames(df)
  ids <- rownames(df)[1:300000]
  fids <- df[ids, "FID"]
  iids <- df[ids, "IID"]
  y <- df[idsT, c(,"FID", "IID", "height")]
```

```

# create data frame with covariates
  cls <- c("FID", "IID", "sex", "age", paste0("pc",1:10))
  covFile <- df[,cls]
  covar <-  paste(c("sex", "age", paste0("pc",1:10)), collapse=" ")
  covFile$sex[covFile$sex==2] <- 0

# select SNPS
  fnSNPs <- "./genotypes/rsids"
  write.table(Glist$study_rsids, row.names = F, col.names = F, quote = F,
              sep = "\t", file = fnSNPs )

# run PLINK
  for(CHR in 1:22){
    tmpOut <- paste0(wd,"LMA_height", "_wbu_ukb_chr",CHR)
    bed <- gsub(".bed", "", Glist$bedfiles)[CHR]
    bim <- Glist$bimfiles[CHR]
    fam <- Glist$famfiles[CHR]

    plink_reg(bedfile=bed, bimfile=bim, famfile=fam, yFile=yFile, fids=fids, ids=ids,
              fnSNPs=fnSNPs, covFile=covFile, covar=covar,  logistic=FALSE,
              wd=wd, fnOut=tmpOut)
  }
```

The two options implemented in qgg are approximations for linear model association (LMA) and mixed linear model association (MLMA). To run a simple regression analysis (*i.e.*, LMA), provide `lma` with the `Glist`, a vector of SNP IDs (`rsids`) and the adjusted phenotype (`y`)

```
  LMA <- lma( y=ytrain[,1], Glist=Glist, rsids=rsids[1:1000], scale=FALSE)
```

To perform the approximation of a mixed linear model analysis three steps are required: 1) build the GRM, 2) estimate the contribution of the GRM to the phenotypic variance using a random effects model , and 3) compute the association statistics that account for this component.

```
  idsT <- train[[1]]
  y <- df[idsT, "height"]
  names(y) <- rownames(df[idsT,])
  X <- model.matrix(~ sex + age + pc1 + pc2 + pc3 + pc4 + pc5 + pc6 + pc7 + pc8 + pc9 + pc10,
                    data=df[idsT,])
  fit <- greml(y=y, X=as.matrix(X), GRMlist=GRMlist, theta=c(0.25,0.05), ncores=12,
               interface="fortran", maxit=10, verbose=TRUE)

  W <- getW(Glist=Glist, ids=idsT, rsids=rsids[1:1000])
  MLMA <- lma(y=y, fit=fit, W = W)
```

Note; for large-scale data sets it may be advantageous to read in chunks of the genotype matrix `W` (for example 10,000 SNPs at a time), and run the analysis in separate steps.

### 6 Gene set enrichment analysis

#### 6.1 Association between genomic features and trait variation

The purpose of gene set enrichment analysis is to identify sets of markers, *i.e.*, genomic features, displaying statistical association with the trait variation, or described in another way, to identify sets of SNPs enriched for causal genetic variants. The main advantages of gene set enrichment analyses are that they (*i*) are computational fast, (*ii*) might not require the genotypes and (*iii*) reduce the issue related to multiple testing by reducing the number of statistical tests performed from the number of genetic variants to the number of genomic features. Gene set enrichment analyses provide a formal statistical modelling framework that allows borrowing and evaluating information obtained across experimental studies.

To evaluate the significance of a test statistic for a given feature set a null hypothesis is needed. Two types of null hypotheses can be used: *competitive* and *self-contained* (Goeman and Bühlmann, 2007; Maciejewski, 2014). The self-contained null hypothesis states that the genomic feature does not display any degree of association with the phenotype, whereas the competitive null hypothesis states that the degree of association of a genomic feature is equal to a similar sized set outside the genomic feature being considered.

For the gene set enrichment methods implemented in qgg we use the competitive null hypothesis. This requires the parameters influencing the summary statistic to be equal to the alternative hypothesis; thus, the number of markers within the true feature set and the sampled sets should be the same, and the correlation structure among markers (due to linkage disequilibrium) should be retained. To fulfil this, an empirical distribution can be obtained using a circular permutation approach (Cabrera et al., 2012). In this approach, the genome was considered to be circular, ordering first the autosomes following any sex chromosomes if included, and ordering the marker test statistics accordingly. The permutation is initiated by randomly selecting one marker element, which then will be the first element in the new rotated genome, and the summary statistic of the first permutation iteration is computed based on the original SNP position. Thus, the markers retain the same order, but receive new marker test statistics. This uncouples any association between the SNPs and feature set, but keeps the correlation structure among test statistics. This permutation approach is then repeated *e.g.* 10,000 times for each genomic feature, and the empirical $p$-values can be obtained through a one-tailed test of the proportion of randomly sampled test statistics that are larger than the observed value.

Currently, four types of gene set enrichment analyses are implemented in qgg. These are breifly described below. For additinal details see Rohde et al. (2016) and Sørensen et al. (2017). The first type is based on counting the number of genetic markers within the feature set that given a significance threshold exhibit association with the trait phenotype,

$$\mathrm{T}_{\mathrm{count}} = \sum_{i=1}^{m_f} \mathbb{I}(|t_i| > t_0), \tag{16}$$

where $m_f$ is the number of markers in a given feature set, $t_i$ is the test statistic for the $i$-th marker (*e.g.* either $t$-statistic or $-\log_{10}(p)$-value), $t_0$ is an arbitrary threshold value for the single marker test statistic, and $\mathbb{I}$ is an indicator function that takes the value one if the argument ($|t_i| > t_0$) is satisfied. Under the null hypothesis, that individually associated markers are distributed randomly, and therefore that the number of associated markers within the feature set is similar to a random set of markers, it can be assumed to follow a hypergeometric distribution, $\mathrm{T}_{\mathrm{count}} \sim \mathrm{Hyper}(m, m_a, m_f)$. That is, $m$ is the total number of markers, $m_a$ is the number of markers that are associated with the trait, and $m_f$ is the number of markers within the feature set.

The second enrichment test is based on the sum of squared test statistics for all markers belonging to a feature set,

$$\mathrm{T}_{\mathrm{sum}} = \sum_{i=1}^{m_f} t_i^2. \tag{17}$$

Here $t_i$ corresponds to the $i$-th test statistic (*i.e.* either $t$-test statistic or marker effect, $\hat{s}_i$) for the $i$-th marker, and $m_f$ is the number of markers located within the feature set. The distribution of $\mathrm{T}_{\mathrm{sum}}$ is unknown; thus, to test the null hypothesis that associated markers are randomly drawn from the total set of markers, an empirical distribution has to be obtained.

The third gene set enrichment approach implemented in qgg is a score-based approach. The approach described here is derived from the first derivative of the likelihood; thus, it is equivalent to the sequence kernel association test (SKAT) (Wu et al., 2011),

$$\mathrm{T}_{\mathrm{score}} = 0.5(\mathbf{y} - \mathbf{Xb})'\mathbf{V}^{-1}\mathbf{G}_f\mathbf{V}^{-1}(\mathbf{y} - \mathbf{Xb}). \tag{18}$$

Here are $\mathbf{b}$ and $\mathbf{V}$ are estimated under the null model, where non-genetic factors are adjusted for, and $\mathbf{G}_f$ is the genomic relationship matrix constructed using those SNPs located within that specific feature set. When the GBLUP model is used as null model it may be advantageous to defined the genetic effects as $\mathbf{g} \sim N(0, \mathbf{G}\sigma_g^2)$, instead of $\mathbf{g}_r \sim N(0, \mathbf{G}_r\sigma_{g_r}^2)$ (where $\mathbf{G}_r$ is the genomic relationship matrix constructed using all the SNPs not located within the feature set), since then each genomic feature will have a specific null model, and thereby the null model has to be fitted for each genomic feature. The test statistic for the score approach can be re-written as:

$$\mathrm{T}_{\mathrm{score}} = 0.5(\hat{\mathbf{e}}'\mathbf{Z}\mathbf{G}_f\mathbf{Z}'\hat{\mathbf{e}}) = 0.5(\hat{\mathbf{e}}'\mathbf{Z}\frac{\mathbf{W}_f\mathbf{W}_f'}{m_f}\mathbf{Z}'\hat{\mathbf{e}}). \tag{19}$$

where $\hat{\mathbf{e}} = \hat{\mathbf{V}}^{-1}(\mathbf{y} - \mathbf{X}\hat{\mathbf{b}})$. For the score approach an empirical distribution under the competitive null hypothesis can be obtained by randomly sampling $m_f$ columns in $\mathbf{W}$.

The fourth, and final type of gene set enrichment approach is the covariance association test (CVAT) (Rohde et al., 2016). The CVAT summary statistic considers the covariance between the total genetic effect (based on all markers, $\hat{\mathbf{g}} = \sum_{i=1}^{m} \mathbf{w}_i\hat{s}_i$), and the feature genetic effects (based on a subset of markers, $\hat{\mathbf{g}}_f = \sum_{i=1}^{m_f} \mathbf{w}_i\hat{s}_i$),

$$\mathrm{T}_{\mathrm{CVAT}} = \mathbf{g}'\mathbf{g}_f = (\mathbf{g}_f' + \mathbf{g}_r')\mathbf{g}_f = \mathbf{g}_f'\mathbf{g}_f + \mathbf{g}_r'\mathbf{g}_f, \tag{20}$$

where $\mathbf{g}_r$ is the genetic effect captured by the markers not included in the feature set, $\hat{\mathbf{g}}_r = \sum_{i=1}^{m_r} \mathbf{w}_i\hat{s}_i$.

#### 6.2 Feature association with gsea

The four types of gene set enrichment analyses described above are all implemented in qgg in the function `gsea`. Below we demonstrate these approaches using feature sets defined by GO terms and KEGG pathways.

First we make sure that feature sets only contain the SNPs that we have in our data, and we restrict the feature sets to those that contain 200+ SNPs.

```
# load SNP association results
  load("./results/LMA_standing_height.Rdata")
  > head(ma)
                      s    stat          p
  rs116720794  0.16190   3.572 0.0003537
  rs3131972   -0.05399  -2.335 0.0195300
  rs12184325   0.16440   3.709 0.0002080
  rs3131962   -0.02884  -1.188 0.2349000
  rs114525117  0.14940   3.272 0.0010680
  rs3115850   -0.05311  -2.182 0.0291200

# GO terms
  load(file="./annotation/goSets.Rdata")
  goSets <- lapply(goSets, function(x){ x[x%in%rownames(ma)] })
  goSets <- goSets[sapply(goSets, length)>199]

# KEGG pathways
  load(file="./annotation/keggSets.Rdata")
  keggSets <- lapply(keggSets, function(x){ x[x%in%rownames(ma)] })
  keggSets <- keggSets[sapply(keggSets, length)>199]
```

The $T_{sum}$ and $T_{count}$ use the results from the linear model association:

```
# GSEA on GO terms
  sumGO <- gsea(stat=as.matrix(ma)[,"stat"]**2, method="sum",sets=goSets, ncores=1, np=100000)
  save(sumGO,file="./results/gsea_sum_goSets_wbu_ukb.Rdata")

  hyperGO <- gsea(stat=as.matrix(ma)[,"p"]**2, method="hyperg",sets=goSets, ncores=1,
              threshold = 0.05)
  save(hyperGO,file="./results/gsea_hyperg_goSets_wbu_ukb.Rdata")

# GSEA on KEGG pathways
  sumKEGG <- gsea(stat=as.matrix(ma)[,"stat"]**2, method="sum",sets=keggSets, ncores=1,
                np=100000)
  save(sumKEGG,file="./results/gsea_sum_keggSets_wbu_ukb.Rdata")

  hyperKEGG <- gsea(stat=as.matrix(ma)[,"p"]**2, method="hyperg",sets=keggSets, ncores=1,
                threshold = 0.05)
  save(hyperKEGG,file="./results/gsea_hyperg_keggSets_wbu_ukb.Rdata"
```

To run $T_{score}$ and $T_{cvat}$ we need the results from `greml`:

```
# load greml results and Glist
  load("./results/ukb_greml.Rdata")
  load("./genotypes/Glist.Rdata")

# GSEA on GO terms
  scoreGO <- gsea(Glist=Glist,fit = fit, sets = goSets, nperm = 10000, method="score",ncores=1)
  save(scoreGO,file="./results/gsea_score_goSets_wbu_ukb.Rdata")

  cvatGO <- gsea(Glist=Glist,fit = fit, sets = goSets, nperm = 10000, method="cvat",ncores=1)
  save(cvatGO,file="./results/gsea_score_goSets_wbu_ukb.Rdata")

# GSEA on KEGG pathways
  scoreKEGG <- gsea(Glist=Glist,fit = fit, sets = keggSets, nperm = 10000, method="score",ncores
      =1)
  save(scoreKEGG,file="./results/gsea_score_keggSets_wbu_ukb.Rdata")

  cvatKEGG <- gsea(Glist=Glist,fit = fit, sets = keggSets, nperm = 10000, method="cvat",ncores=1)
  save(cvatKEGG,file="./results/gsea_score_keggSets_wbu_ukb.Rdata")
```

The results are summarised in Figure S5.

### 7 Pruning for linkage disequilibrium

#### 7.1 Correlated SNPs

Depending on the genetic diversity of the species there can be considerable redundancy in the loci because many loci will be in linkage disequilibrium (LD), especially if one impute SNPs from array to full genome. It is therefore common practise to reduce this redundancy by removing SNPs that are in high LD.

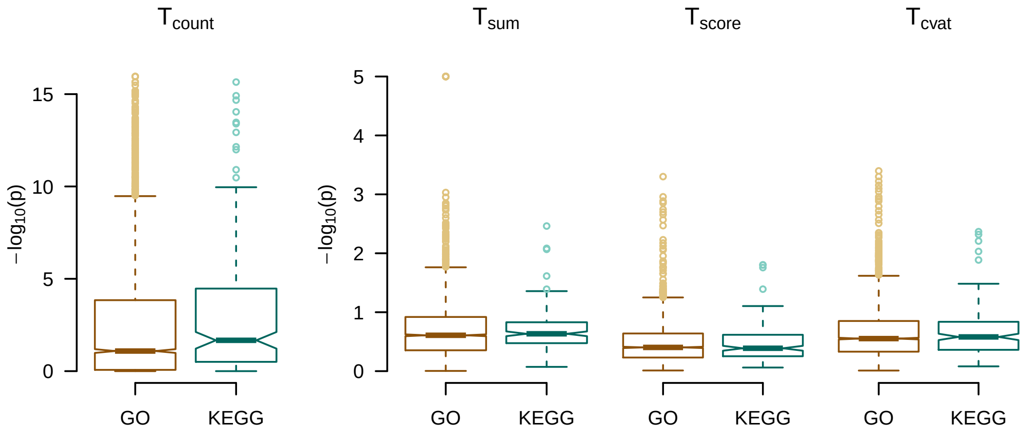

**Figure S5:** Distribution of $-\log_{10}(p)$ for the four types of gene set enrichment analyses implemented in qgg. The gene sets considered were gene ontology (GO) terms and KEGG pathways.

#### 7.2 LD pruning with qgg

The qgg package provides a simple infrastructure to compute LD, and to remove SNPs in high LD within a region of a lead SNP. First, the LD between all SNPs is computed using the `gprep` function with the argument `task="sparseld"`. The LD between pairwise SNPs is computed and stored on the disk as a correlation matrix (∗`.ld`), and the location information is added to the `Glist` object. The LD is computed between all SNPs within a window of size `msize=100`.

```
fnLD <- paste0("./genotypes/UKB_",1:22,".ld")
Glist <- gprep( task="sparseld", Glist=Glist, fnLD=fnLD, msize=100, ncores=1)
save(Glist, file="./genotypes/Glist_wbu_ukb.Rdata")
```

The correlation matrix can be read into `R` using the `getLD` function:

```
ld <- NULL
for(i in 1:22){
  ld[[i]] <- getLD(Glist=Glist, chr=i)
}
```

Then, we can remove all SNPs that are in high LD with lead SNPs (given by the argument `threshold=` in the function `adjLD`) identified from the GWAS.

```
# load files
  load(file="./phenotypes/df_wbu_ukb.Rdata")
  load(file="./genotypes/Glist_wbu_ukb.Rdata")

# run LMA for 30,000 individuals
  df <- df[!is.na(df$standing_height),]
  ids <- rownames(df)
  idsT <- sample(ids, 30000, replace=FALSE)
  y <- df[idsT,"standing_height"]
  names(y) <- rownames(df[idsT,])
  X <- model.matrix(~1, df[idsT,])
  rownames(X) <-rownames(df[idsT,])
  ma <- lma(y = y, X = X, Glist = Glist, ids=idsT, rsids=Glist$study_rsids)

  > dim(ma)
  [1] 599297      4

 # remove SNPs that have an r2 larger than 0.7 with SNPs with p-value of 0.01
  maAdj <- adjLD(stat = ma, Glist = Glist, r2 = 0.7, threshold = 0.01, method = "pruning")

  > dim(maAdj)
  [1] 9257    4
```

### 8 Polygenic risk scores

#### 8.1 Predicting genetic values from SNP effects

In a previous section we described how to predict the genetic values using GBLUP (see Section 4, page 8). An other common approach is to compute polygenic (risk) scores (PRS) as a weighted sum of genotypes,

$$\text{PRS} = \sum_{i=1}^{m} \mathbf{W}_i \hat{\mathbf{s}}_i \tag{21}$$

where $\mathbf{W}_i$ is the $i$-th genotype (allelic counts), and $\hat{\mathbf{s}}_i$ is the weight of the $i$-th genetic variant. The weights can be the log odds ratios (for case-control studies) or the regression coefficients from the single marker analysis (Wray et al., 2007, 2014). The standard approach is to perform the single marker analysis on a training set, and use the estimated SNP effects ($\hat{\mathbf{s}}$) to compute the PRS within the validation population. Typically, the PRS are computed based on markers that have been selected such that the linkage disequilibrium between included markers is below a certain threshold (see Section 7).

#### 8.2 PRS with gscore

The PRS can be computed with the function `gscore`. It is advisable only to include "independent" markers in the prediction; thus, exclude SNPs in high LD (see Section 7). In the example below the `maAdj`-object contains the results from GWAS that has been pruned for LD.

```
# we compute PRS for different p-value thresholds
  threshold <- c(0.00001,0.001,0.01,0.05,0.1,0.2,0.5,0.999)

  for ( thold in threshold) {
        fnGRS <- paste("./results/prs_height_",thold,".Rdata",sep="")
        prs <- gscore(Glist=Glist,stat=maAdj,ncores=1)
        save(prs, file="fnGRS")
  }
```

The accuracy of prediction can then be obtained with the function `acc`,

```
# for one training set
  yt <- na.omit(ytrain[,1])
  idsT <- names(yt)
  idsV <- names(y)[!names(y)%in%idsT]
  yobs <- y[idsV]
  ypred <- prs[idsV,1]
  R2 <- acc(yobs=yobs,ypred=ypred}
```

### 9 Joint estimation of marker effects

#### 9.1 Multi-marker association model

The association analysis between SNP-markers and complex trait phenotypes is typically performed as a SNP-by-SNP analysis; however, multi-marker association models, where the SNP effects are estimated jointly, can be an alternative. Consider the linear mixed model,

$$\mathbf{y} = \mathbf{1}\mu + \mathbf{W}_k \mathbf{s}_k + \mathbf{e}, \tag{22}$$

where $\mathbf{y}$ is the vector of phenotypic observations adjusted for known fixed effects, $\mathbf{1}$ is a vector of ones, $\mu$ is the overall mean, $\mathbf{W}_k$ is the centred and scaled genotype matrix for $k$ markers, $\mathbf{s}$ is the corresponding vector of SNP effects, and $\mathbf{e}$ is the vector of residuals. The SNP effects can be obtained using a direct method,

$$\mathbf{s} = (\mathbf{W}'\mathbf{W} + \mathbf{I}\lambda)^{-1}\mathbf{W}'\mathbf{y}, \tag{23}$$

where $\lambda = \sigma_e^2/\sigma_s^2$. However, the dimension of $(\mathbf{W}'\mathbf{W} + \mathbf{I}\lambda)$ is $m$-by-$m$, thus, when $m$ is large the inversion becomes computationally intensive. Alternatively, this can be solved using an iterative approach, like Gauss-Seidel. In qgg, we have implemented Gauss-Seidel using a so-called Gauss-Seidel with residual update (GSRU) method. Let $\mathbf{w}_i$ denote the $i$-th column vector of the genotype matrix $\mathbf{W}$. Then, we can update the SNP effect of the $i$-th element in $\mathbf{s}$ as:

$$\mathbf{s}_i^{new} = \frac{\mathbf{w}_i'(\mathbf{y} - \mathbf{W}_{-i}\mathbf{s}_{-i})}{\mathbf{w}_i'\mathbf{w}_i + \lambda} \tag{24}$$

The term, $\mathbf{y} - \mathbf{W}_{-i}\mathbf{s}_{-i}$, corresponds to the residual corrected for all effects, except the $i$-th SNP ($s_i$).

$$\mathbf{s}_i^{new} = \frac{\mathbf{w}_i'(\mathbf{y} - \mathbf{W}_{-i}\mathbf{s}_{-i} - \mathbf{w}_i s_i + \mathbf{w}_i s_i)}{\mathbf{w}_i'\mathbf{w}_i + \lambda} \quad (25)$$

$$= \frac{\mathbf{w}_i'(\mathbf{y} - \mathbf{W}\mathbf{s} + \mathbf{w}_i s_i)}{\mathbf{w}_i'\mathbf{w}_i + \lambda} \quad (26)$$

$$= \frac{\mathbf{w}_i'(\mathbf{e} + \mathbf{w}_i s_i)}{\mathbf{w}_i'\mathbf{w}_i + \lambda} \quad (27)$$

$$= \frac{\mathbf{w}_i'\mathbf{e} + \mathbf{w}_i'\mathbf{w}_i s_i}{\mathbf{w}_i'\mathbf{w}_i + \lambda} \quad (28)$$

$$= \frac{\mathbf{w}_i'\mathbf{e}^{old} + \mathbf{w}_i'\mathbf{w}_i s_i}{\mathbf{w}_i'\mathbf{w}_i + \lambda} \quad (29)$$

(30)

Updating the residuals $\mathbf{e}$ is necessary after computation of each effect. The updating after each effect is:

$$\mathbf{e}^{new} = \mathbf{e}^{old} - \mathbf{w}_i'(s_i^{new} - s_i) \quad (31)$$

Thus, Equation 30 and 31 describes one iteration in the Gauss-Seidel Residual Update algorithm for the $i$-th genetic marker.

#### 9.2 Joint marker estimation with gsolve

In qgg the multi-marker association approach is conducted with the function `gsolve`.

```
fit <- gsolve( y=y, X=X, Glist=Glist, rsids=rsids, lambda=lambda, maxit=10, tol=1e-5 )
```

The `fit`-object contains the re-estimated SNP effects (`fit$s`), and the predicted genetic effects (`fit$g`), which is computed as in Equation 21.

In Figure S6 we compare the variance explained ($R^2$) in the validation set from predictions using SNP effects estimated with a standard linear model association approach (here PLINK (Purcell et al., 2007)) to predictions using SNP effects jointly estimated using the Gauss-Seidel Residual Update algorithm.

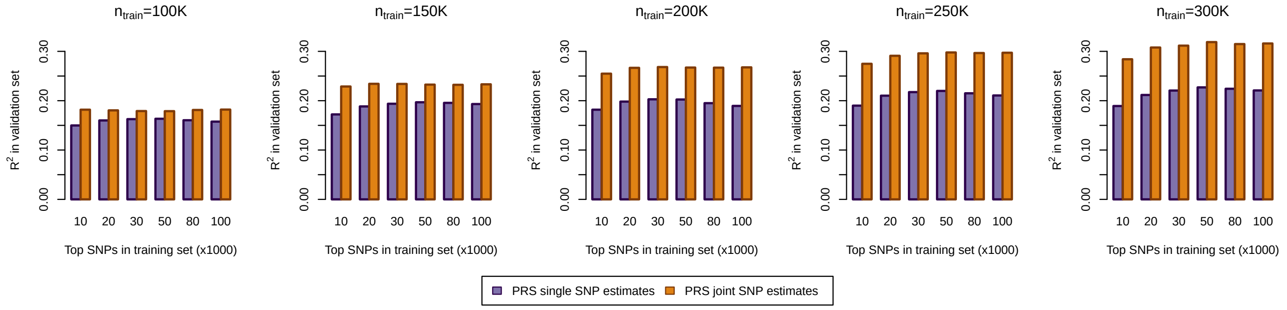

**Figure S6:** Comparison of variance explained ($R^2$) in the validation set from prediction (PRS) using SNP effects estimated using a standard linear model association approach (PLINK) and SNP effects joint estimated using the Gauss-Seidel Residual Update algorithm. The prediction models were constructed using top 10K, 20K, 30K, 50K, 80K or 100K associated SNPs in the training set (we used a range of different training set sizes, $n_{train}$).

Prediction variance explained for human height lies typically between 0.15-0.4 (Ge et al., 2019; Lloyd-Jones et al., 2019; Lello et al., 2018); but largely depend on training size, and if the training/validation set contains related individuals. Lello et al. (2018) reported an $R^2 \sim 0.4$ for human height, but their data set contained related indivudals which will increase the accuracy. Using the Gauss-Seidel Residual Update algorithm to jointly estimate the SNP effects in the training set we got a maximum variance explained of 0.32, which is equally high to comparable prediction schemes.

### 10 Computational times

The qgg package utilises multi-core processing with openMP and multithreaded matrix operations implemented in BLAS libraries (OpenBLAS, ATLAS or MKL) which provides an efficient computational infrastructure for large-scale genetic data.

The GRM is computed using distributed matrix multiplication (multi-core and multi-node), estimation of variance components is based on an average information restricted maximum likelihood (AI-REML) (currently multi-core within node) and marker test statistics are computed independently using distributed computing (multi-core and multi-node). The computation time can be broken down into three steps: (1) building the GRM ($O(MN^2)$), (2) estimating variance components ($O(N^3)$) and (3) computing association statistics for each marker ($O(MN^2)$).

In the following we illustrate these features on the construction of the GRM and on estimation of variance components. The GRM is computed using distributed matrix multiplication (multi-core and multi-node), estimation of variance components is based on an average information restricted maximum likelihood (AI-REML) (currently multi-core within node). To illustrate the computation time using multi-core processing we constructed the GRM using $n = 50000$ individuals with $m = \{10000, 50000, 100000, 250000, 500000\}$ SNP markers using 1, 4, 8 or 16 cores. Similarily, we estimated variance components using $n = \{1000, 5000, 10000, 20000, 50000\}$ individuals, and the GRM constructed from 100000 markers. All analyses were performed using R (version 3.5.0) on the GenomeDK HPC cluster at Aarhus University, Denmark (http://genome.au.dk), with 60 GB memory allocated per computer node. The results are summarised in Figure S7.

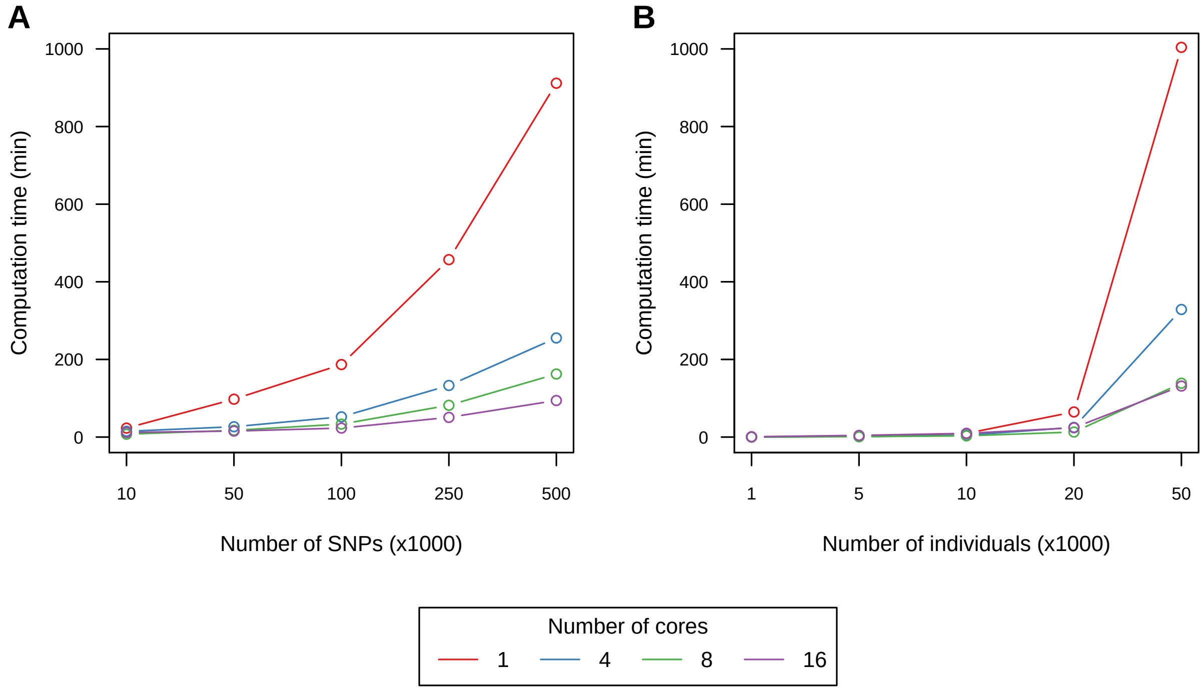

**Figure S7:** Computation times for **A**) construction of GRM for 50K individuals, and **B**) estimating variance components.

### Appendix: R-code for UKB example

The appendix contains the R-code used to produce the results presented within the main manuscript.

#### Prepare phenotype data for qgg

```
#------------------------------------------------------------------------------------------------#
# after the UKB phenotype data have been decrypted with the supplied UKB programs,
# the R package ukbtools can be used to easily create a dataframe within R.

# install and load packages
  install.packages("ukbtools")
  library(ukbtools)

  df <- ukb_df("ukbxxxx", path="./ukb_rawfiles/")
  save(df, file="./phenotypes/ukb_data.Rdata")

#------------------------------------------------------------------------------------------------#
# restrict to the White-British cohort, and remove related individuals, individuals with
    chromosomal aneuploidy, and if an individual have a genotype missing rate above 0.05.

# extract the White British cohort
  is.British <- df[, "ethnic_background_0_0" ]=="British"
  df <- df[is.British,]
  is.white <- df[,"genetic_ethnic_grouping_0_0"]=="Caucasian"
  is.white[is.na(is.white)] <- FALSE
  df <- df[is.white,]

# remove related individuals
  rel <- read.table(file="./genotypes/ukbxxxx_rel_syyyyyy.dat",sep=" ", header=TRUE)
  ids1 <- rel[,"ID1"]
  ids2 <- rel[,"ID2"]
  ids_related <- ids2[!ids2%in%ids1]

  is.related <- df[,"eid"]%in%ids_related
  df <- df[!is.related,]

# remove individuals with aneuploidy
  is.aneuploidy <- df[,"sex_chromosome_aneuploidy_0_0"]=="Yes"
  is.aneuploidy[is.na(is.aneuploidy)] <- FALSE
  df <- df[!is.aneuploidy,]

# remove individuals with genotype missingness above 0.05
  is.missing <- df[,"missingness_0_0"]<=0.05
  df <- df[is.missing,]

#------------------------------------------------------------------------------------------------#
# manipulate the dataframe for downstream analyses
  df$eid <- as.character(df$eid)
  rownames(df) <- df$eid

# select coloums to be included in the new dataframe
  cls <- c("sex_0_0",
           "year_of_birth_0_0",
           "genetic_principal_components_0_1",
           "genetic_principal_components_0_2",
           "genetic_principal_components_0_3",
           "genetic_principal_components_0_4",
           "genetic_principal_components_0_5",
           "genetic_principal_components_0_6",
           "genetic_principal_components_0_7",
           "genetic_principal_components_0_8",
           "genetic_principal_components_0_9",
           "genetic_principal_components_0_10",
           "standing_height_0_0")

  df <- df[,cls]
  colnames(df) <- c("sex","age",paste("pc",1:10,sep=""), "height")
  df[,"age"] <- 2018-df[,"age"]

# centre and scale the phenotype (mean=0, sd=1)
  df$height <- scale(df$height)
  save(df,file="./phenotypes/df_wbu_ukb.Rdata")
```

```

# adjust y (human standing height) for sex, age and genetic principal components
  df$height_scaled <- NA
  isNA <- is.na(df[,"height"])
  X <- model.matrix( ~ sex + age + pc1 + pc2 + pc3 + pc4 + pc5 + pc6 + pc7 + pc8 + pc9 + pc10,
      data=df[!isNA,])
  fit <- fastlm(y=df[!isNA,"height"],X=X)
  df[!isNA,"height_scaled"] <- df[!isNA,"height"]-fit$yhat
  save(df,file="./phenotypes/df_wbu_ukb_scaled.Rdata")

#----------------------------------------------------------------------------------------------#
# setup training-validation sets with different sizes
  noNA <- !is.na(df$height_scaled)
  idsT <- rownames(df)[noNA]

  ntrain <- rep(c(100000,150000,200000,250000,300000), each=5)
  train <- sapply(ntrain, function(x){sample(idsT,x)})

  ytrain <- NULL
  for (i in 1:length(train)) {
        yobs <- df$height_scaled
        names(yobs) <- rownames(df)
        isT <- names(yobs)%in%train[[i]]
        yobs[!isT] <- NA
        ytrain <- cbind(ytrain,as.matrix(yobs))
  }

  reps <- as.character(rep(rep(1:5),times=5))
  colnames(ytrain) <- paste(rep(c("100K","150K","200K","250K","300K"),each=5),reps,sep="_")

  save(ytrain,file="./phenotypes/ytrain_scaled_wbu_ukb.Rdata")
  save(train,file="./phenotypes/train_wbu_ukb.Rdata")
```

#### Prepare genotype data for qgg

To prepare the genotype file it is required that the UKB binary PLINK (Purcell et al., 2007) files have been downloaded and decrypted as specified by the UK Biobank Resource (http://biobank.ndph.ox.ac.uk/showcase/download.cgi).

```
#----------------------------------------------------------------------------------------------#
# prepare input parameters to prepare the Glist
  library(qgg)

  load(file="./phenotypes/df_wbu_ukb.Rdata")
  idsWBU <- rownames(df)

# specify the path and names of the binary PLINK files
  bimfiles <- paste("./genotypes/ukb_snp_chr",1:22,"_v2.bim",sep="")
  bedfiles <- paste("./genotypes/ukb_cal_chr",1:22,"_v2.bed",sep="")
  famfiles <- paste("./genotypes/ukbxxxxxx_cal_chr",1:22,"_v2_syyyyyy.fam",sep="")

# specify which individuals should be used in project (this is optional - if left out all
    individuals in fam files are used)
  ids <- idsWBU

# name of genotype file used in all qgg analyses - remember *.raw extension
  fnRAW <- "./genotypes/wbu_ukb.raw"

# construct Glist
  Glist <- gprep(study="WBU_UKB", fnRAW=fnRAW, bedfiles=bedfiles, bimfiles=bimfiles,
                 famfiles=famfiles, ids=ids, ncores=1)

# specify which SNPs that should be used in the analyses
  isMAF <- Glist$maf <=0.01
  isMISS <- Glist$nmiss/length(Glist$study_ids) > 0.05
  isMHC <- Glist$chr==6 & Glist$position > 25602429 & Glist$position < 33471466
  delete <- isMAF | isMISS | isMHC
  Glist$study_rsids <- Glist$rsids[!delete]

  save(Glist, file="./genotypes/Glist_wbu_ukb.Rdata", compress=FALSE)
```

#### REML analysis

Below we estimate the heritability for human height for $n = 10K, 20K, 30K, 50K$, and partition the total genetic variance to each of the 22 autosomal chromosomes using $n = 50K$.

```
#-----------------------------------------------------------------------------------------------#
# load the Glist, and prepare the phenotype and covariate data for greml
  library(qgg)

  load(file="/genotypes/Glist_wbu_ukb.Rdata")
  load(file="/phenotypes/df_wbu_ukb.Rdata")
  y <- na.omit(as.matrix(df[,"standing_height",drop=FALSE])[,1])
  X <- model.matrix(~ sex + age + pc1 + pc2 + pc3 + pc4 + pc5 + pc6 + pc7 + pc8 + pc9 + pc10,
                    data=df[names(y),])

#-----------------------------------------------------------------------------------------------#
# estimate heritability using different study population sizes.

# setup input parameters
  ncores <- 12
  nrep <- 5
  ntrain <- c(10000,20000,30000,50000)

# construct GRM and estimate variance components for the different training population sizes.
  for ( i in c(1:nrep) ) {
        idsG <- sample(names(y),max(ntrain))
        fnG <- "./grm/G.grm"
        GRMlist <- grm(Glist=Glist, ids=idsG, rsids=Glist$study_rsids, scale=TRUE, msize=100,
                       ncores=ncores, fnG=fnG, overwrite=TRUE)
        fit <- vector(length=length(ntrain),mode="list")

        for ( j in 1:length(ntrain) ) {
              idsT <- idsG[1:ntrain[j]]
              fit[[j]] <- greml(y=y[idsT], X=as.matrix(X[idsT,]), GRMlist=GRMlist,
                                theta=c(0.25,0.05), ncores=ncores, interface="fortran", maxit=10,
                                verbose=TRUE)
        }
        fnGREML <- paste("./results/greml_rep",i,"_100K_wbu_ukb.Rdata",sep="")
        save(fit, file=fnGREML)
  }

#-----------------------------------------------------------------------------------------------#
# partition of genetic variance across autosomal chromosomes.

# setup input parameters
  nrep <- 5
  ntrain <- 50000

  fit <- vector(length=nrep,mode="list")

# construct GRM for each autosomal chromosome and fit all 22 genetic components in one GREML
  for ( i in 1:nrep) {
        idsG <- sample(names(y),ntrain)
        ifnG <- paste("./grm/G",1:22,".grm",sep="")
        GRMlist <- NULL

        for ( chr in 1:22 ) {
              rsidsChr <- Glist$rsids[Glist$chr==chr]
              rsidsChr <- rsidsChr[rsidsChr%in%rsids]
              GRMlist[[chr]] <- grm(Glist=Glist, ids=idsG, rsids=rsidsChr, scale=TRUE,
                                    msize=100, ncores=16, fnG=fnG[chr], overwrite=TRUE)
        }

        names(GRMlist)<- paste("CHR",1:22,sep="")
        GRMlist <- mergeGRM( GRMlist )

        fit[[i]] <- greml(y=y[idsG], X=as.matrix(X[idsG,]), GRMlist=GRMlist, theta=rep(0.01,23),
                          ncores=16, interface="fortran", maxit=20, verbose=TRUE)

        save( fit, file="./results/greml_chr_50K_wbu_ukb.Rdata")
  }
```

#### Single marker analysis

Then we estimate the SNP effects using linear model association. The SNP effects are used later for gene set enrichment analysis, and to construct the polygenic risk scores.

```
#-----------------------------------------------------------------------------------------------#
# estimate SNP effects with lma
  library(qgg)

# load scaled phenotypes for the training populations
  load(file="./phenotypes/ytrain.Rdata")

# run analysis
  for( i in 1:ncol(ytrain) ){
       ma <- lma( y=ytrain, Glist=Glist, rsids=Glist$study_rsids, scale=FALSE)
       save(ma, file=paste0("./results/ma_wbu_ukb_", colnames(ytrain)[i] ,".Rdata"))
  }
```

#### Prune SNP markers for LD

Before we construct the polygenic risk scores we compute the LD between SNP-pairs, and remove SNPs that are in high LD ($r^2 < 0.7$).

```
#-----------------------------------------------------------------------------------------------#
# compute LD - this is done with the gprep function

  library(qgg)

  load(file="./genotypes/Glist_wbu_ukb.Rdata")
  fnLD <- paste0("./genotypes/LDmat_wbu_ukb",1:22,".ld")
  Glist <- gprep( task="sparseld", Glist=Glist, fnLD=fnLD, msize=100, ncores=6)
  save(Glist, file="./genotypes/Glist_wbu_ukb.Rdata", compress=FALSE)

#-----------------------------------------------------------------------------------------------#
# adjust summary statistics from lma

  for( i in 1:ncol(ytrain) ){
       load(file=paste0("./results/ma_wbu_ukb_", colnames(ytrain)[i] ,".Rdata"))
       maAdj <- adjLD(stat = ma, Glist = Glist, r2 = 0.7, threshold = 0.01, method = "pruning")
       save(maAdj, file=paste0("./results/maAdj_wbu_ukb_", colnames(ytrain)[i] ,".Rdata"))
  }
```

#### Polygenic risk scores for human height

We will construct polygenic risk scores using a range of $p$-value thresholds, but also using top-X associated SNPs (we will use the latter results to compare with the results from the predictions using the jointly estimated SNP effects).

```
#-----------------------------------------------------------------------------------------------#
# PRS for p-value threshold
  load(file="./genotypes/Glist_wbu_ukb.Rdata")
  load(file="./phenotypes/ytrain.Rdata")
  cnames <- colnames(ytrain)

  threshold <- c(0.00001,0.001,0.01,0.05,0.1,0.2,0.5,0.999)

  for( thold in threshold) {

        for( i in 1:ncol(ytrain)){
             fnPRS <- paste0("./results/prs_", cnames[i],"_thr" ,thold,"_R70_wbu_ukb.Rdata")
             load(paste0("./results/maAdj_wbu_ukb_", colnames(ytrain)[i] ,".Rdata"))
             ma <- ma[ma$p<threshold[thold],]
             prs <- gscore(Glist=Glist, stat=ma, ncores=8)
             save(prs,file=fnPRS)
        }
  }

#-----------------------------------------------------------------------------------------------#
# PRS for top SNPs
  load(file="./genotypes/Glist_wbu_ukb.Rdata")
  load(file="./phenotypes/ytrain.Rdata")
```

```
  cnames <- colnames(ytrain)

  top <- c(10000,20000,30000,50000,80000,100000)

     for( i in 1:length(top)) {

          for( i in 1:ncol(ytrain)){
               fnPRS <- paste0("./results/prs_cnames[i],"_top",top[i],"_R70_wbu_ukb.Rdata")
               load(paste0("./results/maAdj_wbu_ukb_", cnames(ytrain)[i] ,".Rdata"))
               ma <- ma[order(ma$p, decreasing=FALSE),]
               ma <- ma[1:top[i],]
               prs <- gscore(Glist=Glist, stat=ma, ncores=8)
               save(prs,file=fnPRS)
          }
  }
```

The final step is then to compute the prediction accuracies within the validation sets.

```
#--------------------------------------------------------------------------------------------#
# accuracy for PRS with p threshold
  load(file="./phenotypes/ytrain.Rdata")
  cnames <- colnames(ytrain)

  accuracy <- NULL
  threshold <- c(0.00001,0.001,0.01,0.05,0.1,0.2,0.5,0.999)

  for( thold in threshold){

        for( i in 1:ncol(ytrain)){
             load(paste0("./results/prs_", cnames[i],"_thr", thold, "_R70_wbu_ukb.Rdata"))
             idsT <- rownames(ytrain[!is.na(ytrain[,i])])
             idsV <- rownames(ytrain[is.na(ytrain[,i])])

             yobs <- ytrain[!is.na(ytrain[,i]),i]
             ypred <- prs[idsV,]

             reps <- substr(cnames[i],nchar(cnames[i]),nchar(cnames[i]))
             ntrain <- gsub("\\_.*","",gsub("standing_height_", "",cnames[i]))
             accuracy <- rbind(accuracy,c(thold,ntrain,reps,acc(yobs=yobs,ypred=ypred)))
        }
  }
  save(accuracy, file="./results/accuracy_prs_threshold_R70_wbu_ukb.Rdata")

#--------------------------------------------------------------------------------------------#
# accuracy for PRS with top SNP
  accuracy <- NULL
  top <- c(10000,20000,30000,50000,80000,100000)

  for( tp in top){

        for( i in 1:ncol(ytrain)){
             load(paste0("./results/prs_", cnames[i],"_top", tp, "_R70_wbu_ukb.Rdata"))
             idsT <- rownames(ytrain[!is.na(ytrain[,i])])
             idsV <- rownames(ytrain[is.na(ytrain[,i])])

             yobs <- ytrain[!is.na(ytrain[,i]),i]
             ypred <- prs[idsV,]

             reps <- substr(cnames[i],nchar(cnames[i]),nchar(cnames[i]))
             ntrain <- gsub("\\_.*","",gsub("standing_height_", "",cnames[i]))
             accuracy <- rbind(accuracy,c(tp,ntrain,reps,acc(yobs=yobs,ypred=ypred)))
        }
  }
    save(accuracy, file="./results/accuracy_prs_top_R70_wbu_ukb.Rdata")
```

#### Joint estimation of SNP effects using Gauss-Seidel residual update algorithm

Now we re-estimate the SNP effects of the top associated SNPs using the Gauss-Seidel algorithm .

```
#--------------------------------------------------------------------------------------------#
# joint estimation of marker effects
  library(qgg)

  load(file="./genotypes/Glist_wbu_ukb.Rdata")
```

```
load(file="./phenotypes/ytrain.Rdata")
cnames <- colnames(ytrain)

top <- c(10000,20000,30000,50000,80000,100000)
h2 <- 0.7

for( tp in top){
    g <- matrix(NA,nrow=Glist$n,ncol=ncol(ytrain))
    rownames(g) <- Glist$ids
    colnames(g) <- colnames(ytrain)

    for( i in 1:ncol(ytrain)){
         load(file=paste0("./results/ma_WBU_UKB_", colnames(ytrain)[i] ,".Rdata"))
         fnG <- paste0("/results/ghat_top",top,"_wbu_ukb.Rdata")

         rsids <- rownames(ma$p)[order(ma$p, decreasing=FALSE)][1:top]
         yt <- na.omit(ytrain[,i])
         Xt <- model.matrix(yt~1)
         lambda <- length(rsids)*(1-h2)/h2

         fit <- gsolve(y=yt, X=Xt, Glist=Glist, rsids=rsids, lambda=lambda,
                       maxit=10, tol=1e-5)
         g[,i] <- fit$g
    }
    save(g, file=fnG)
}
```

The accuracies of prediction in the validation set is then computed.

```
#---------------------------------------------------------------------------------------------#
# accuracy for Gauss-Seidel PRS
  accuracy <- NULL
  top <- c(10000,20000,30000,50000,80000,100000)

  for( tp in top){
     load(file=paste("./results/ghat_top",top,"_wbu_ukb.Rdata",sep=""))

     for( i in 1:ncol(ytrain)){
             idsT <- rownames(ytrain[!is.na(ytrain[,i])])
             idsV <- rownames(ytrain[is.na(ytrain[,i])])

             yobs <- ytrain[!is.na(ytrain[,i]),i]
             ypred <- g[idsV,i]

             reps <- substr(cnames[i],nchar(cnames[i]),nchar(cnames[i]))
             ntrain <- gsub("\\_.*","",gsub("standing_height_", "",cnames[i]))
             accuracy <- rbind(accuracy,c(tp,ntrain,reps,acc(yobs=yobs,ypred=ypred)))
       }
  }
```

#### Gene set enrichment analysis

Finally, we perform a gene set enrichment analysis using genomic features defined by gene ontology terms. Here we used the $T_{sum}$ approach.

```
#---------------------------------------------------------------------------------------------#
# GSEA using GO terms
  library(qgg)

  load(file="./annotation/goSets.Rdata")
  load(file="./genotypes/Glist_wbu_ukb.Rdata")

# make sure only rsids that are in the data set are in the gene sets
  goSets <- lapply(Glist, function(x){x[x%in%Glist$study_rsids]})

# we use the LMA results performed on 300K individuals
  cls <- paste0("standing_height_300K_",1:5)

  res <- NULL
  for( i in cls ){
       load(file=paste0("./results/ma_WBU_UKB_", cls, ".Rdata"))
       res[[i]] <- gsea(stat = as.matrix(ma)[Glist$study_rsids,"stat"]**2, sets = goSets,
                        method = "sum", nperm = 100000)
```

```
}
save(res, file="./results/gsea_sum_standing_height_300K_wbu_ukb.Rdata")
```
